# Insight and Inference for DVARS

**DOI:** 10.1101/125021

**Authors:** Soroosh Afyouni, Thomas E. Nichols

## Abstract

Estimates of functional connectivity using resting state functional Magnetic Resonance Imaging (rs-fMRI) are acutely sensitive to artifacts and large scale nuisance variation. As a result much effort is dedicated to preprocessing rs-fMRI data and using diagnostic measures to identify bad scans. One such diagnostic measure is DVARS, the spatial standard deviation of the data after temporal differencing. A limitation of DVARS however is the lack of concrete interpretation of the absolute values of DVARS, and finding a threshold to distinguish bad scans from good. In this work we describe a variance decomposition of the entire 4D dataset that shows DVARS to be just one of three sources of variation we refer to as *D*-var (closely linked to DVARS), *S*-var and *E*-var. *D*-var and *S*-var partition the variance at adjacent time points, while *E*-var accounts for edge effects; each can be used to make spatial and temporal summary diagnostic measures. Extending the partitioning to global (and non-global) signal leads to a rs-fMRI DSE ANOVA table, which decomposes the total and global variability into fast (*D*-var), slow (*S*-var) and edge (*E*-var) components. We find expected values for each component under nominal models, showing how *D*-var (and thus DVARS) scales with overall variability and is diminished by temporal autocorrelation. Finally we propose a null sampling distribution for DVARS-squared and robust methods to estimate this null model, allowing computation of DVARS p-values. We propose that these diagnostic time series, images, p-values and ANOVA table will provide a succinct summary of the quality of a rs-fMRI dataset that will support comparisons of datasets over preprocessing steps and between subjects.

## 1. Introduction

Functional connectivity obtained with resting state functional magnetic resonance imaging (rs-fMRI) is typically computed by correlation coefficients between different brain regions, or with a multivariate decomposition like Independent Components Analysis (Cole et al., 2010). Both approaches can be corrupted by artifacts due to head motion or physiological effects, and much effort is dedicated to preprocessing rs-fMRI data and using diagnostic measure to identify bad scans.

Smyser et al. (2011) proposed and Power et al. (2012) popularized a measure to characterize the quality of fMRI data, an image-wide summary that produces a time series that can detect problem scans. They called their measure DVARS, defined as the spatial standard deviation of successive difference images. In fact, DVARS can be linked to old statistical methods developed to estimate noise variance in the presence of drift (see Appendix A for DVARS history).

While DVARS appears to perform well at the task of detecting bad scans — bad pairs of scans — it does not have any absolute units nor a reference null distribution from which to obtain p-values. In particular, the typical “good” values of DVARS varies over sites and protocols which makes it difficult to create comparable summaries of data quality across data sets. The emergence of the large scale data sets such as the Human Connectome Project (HCP) (>1k subjects) and the UK Biobank (>10k subjects) further motivates the need for automated, yet reliable, quantitative techniques to control and improve the data quality.

The purpose of this work is to provide a formal description of DVARS as part of a variance decomposition of the data, propose more interpretable standardized versions of DVARS, and compute DVARS p-values for a null hypothesis of homogeneity.

The remainder of this work is organized as follows. We first describe the variance decomposition for the 4D data and how this relates to traditional DVARS, and other new diagnostic measures it suggests. Then we describe a sampling distribution for DVARS under the null hypothesis, and mechanisms for estimating the parameters of this null distribution. We establish the validity and sensitivity of the DVARS test with simulations, and use two different fRMI cohorts to demonstrate how both the DVARS test and our‘DSE’ decomposition are useful to identify problem subjects and diagnose the source of artifacts within individual subjects.

## 2. Theory

Here we state our results concisely relegating full derivations to Appendices.

### 2.1. Notation

For *T* time-points and *I* voxels, let the original raw rs-fMRI data at voxel *i* and *t* be 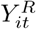. Denote the mean at voxel *i* as 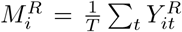, and by *m*^*R*^ some type of overall mean value (i.e. a summary of the mean image 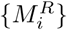, like median or mean). We take as our starting point for all calculations the centered and scaled data:

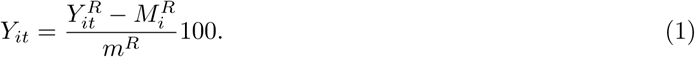

The scaling ensures that typical brain values before centering are around 100 and are comparable across datasets; centering allows mean squared computations to be interpreted as variance.

### 2.2. DSE Variance Decomposition

Denote the total (“all”) variance at scan t as

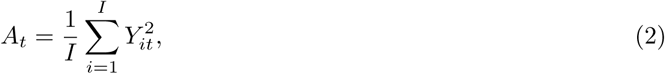

which can also be seen as the average of voxel-wise variances at scan *t*. Define two mean squared terms, one for fast (“differenced”) variability

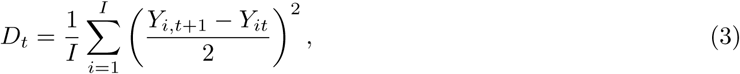

the half difference between time *t* and *t*+1 at each voxel, squared and averaged over space, and one for slow variability

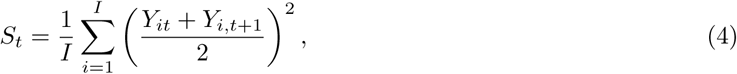

the average between *t* and *t* + 1 at each voxel, squared and averaged over space.

We then have the following decomposition of the average variance at time points *t* and 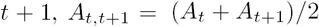

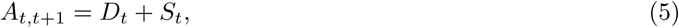

for *t* = 1,…, *T* −1. This has a particularly intuitive graphical interpretation: If we plot *D*_*t*_ and *S*_*t*_ at *t*+1/2, they sum to the midpoint between *A*_*t*_ and *A*_*t*+1_ found at *t* + 1/2 (see Fig. 1). Since

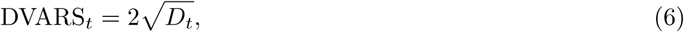

we now have a concrete interpretation for DVARS, with 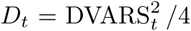 being the “fast” SS component in the average variance at *t* and *t* + 1.

**Figure 1:**
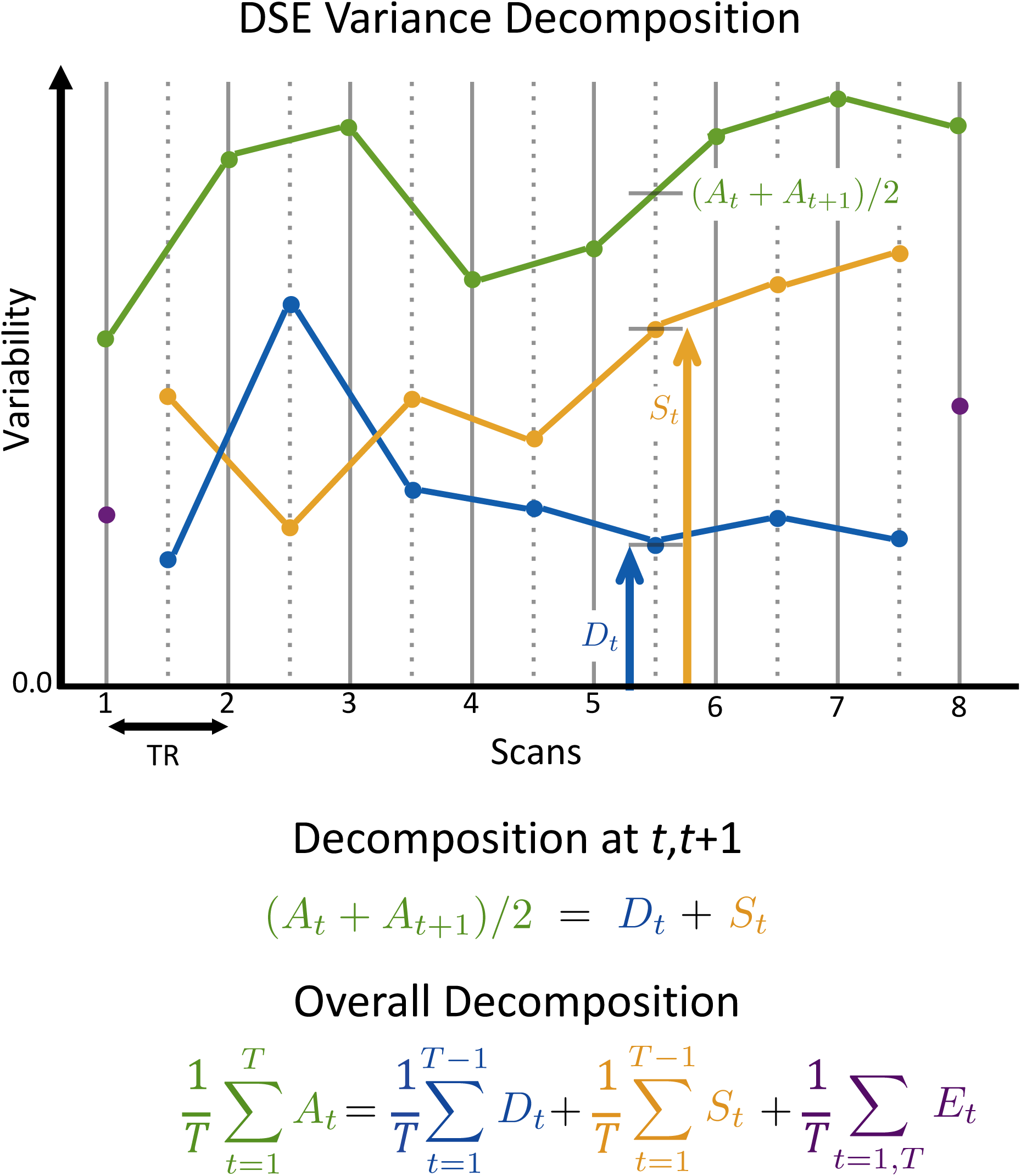
Illustration of the DSE decomposition, where *A*_*t*_ (green) is the total sum-of-squares at each scan, *D*_*t*_ (blue) is the sum-of-squares of the half difference of adjacent scans, *S*_*t*_ (yellow) is the sum-of-squares of the average of adjacent scans, and *E*_*t*_ is the edge sum-of-squares at times 1 and 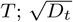 is proportional to DVARS. The *D* and *S* components for index *t* (*D*_*t*_ and *S*_*t*_) sum to *A* averaged between *t* and *t* + 1 ((*A*_*t*_ + *A*_*t*+1_)/2). Note how the *S* and *D* time series allow insight to the behavior of the total sum-of-squares: The excursion of *A* around *t =* 2, 3 arise from fast variance while the rise for *t* ≥ 6 is due to slow variance. For perfectly clean, i.e. independent data, *D* and *S* will converge and each explain approximately half of *A.*

This also leads to a decomposition of the total variance over all scans: With averages *A*, *D*, *S* and *E* defined in Table 1 (row 1) we have the following “DSE” decomposition

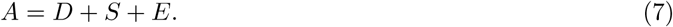

That is, the total variance (“*A*-var”) in the 4D dataset is the sum of terms attributable to fast (“*D*-var”), slow (“*S*-var”) and edge variability (“*E*-var”). *D* is also 1/4 the average mean squared difference (MSSD; see Appendix A). Each term in the “DSE” decomposition can be split into global and non-global components, as shown in Table 1, rows 2-3 (as also noted by Burgess et al. (2016) for *D*_*t*_).

**Table 1:** Make up of the DSE ANOVA table giving a mean squared (MS) variance decompositions of resting-state fMRI data. The first row shows how the total MS can be split into 3 terms, in the second through 4th columns, *A* = *D* + *S* + *E*. The first column likewise shows how total MS can be decomposed in to that explained by a spatially global time series (second row) and a non-global or residual-global component (third row), *A* = *A*_*G*_ + *A*_*N*_. Likewise, each row and column sums accordingly: *A*_*G*_ = *D*_*G*_ + *S*_*G*_ + *E_G_, D* = *D*_*G*_ + *D*_*N*_, etc. Terms are shown here as MS for brevity, but are best reported in root mean squared (RMS) units. See Table 2 for definitions of the time series variables.

|  | A-var | D-var | S-var | E-var |
| --- | --- | --- | --- | --- |
| Whole | $A = \frac{1}{T} \sum_{t=1}^T A_t$ | $D = \frac{1}{T} \sum_{t=1}^{T-1} D_t$ | $S = \frac{1}{T} \sum_{t=1}^{T-1} S_t$ | $E = \frac{1}{T} \sum_{t=1, T} E_t$ |
| Global | $A_G = \frac{1}{T} \sum_{t=1}^T A_{Gt}$ | $D_G = \frac{1}{T} \sum_{t=1}^{T-1} D_{Gt}$ | $S_G = \frac{1}{T} \sum_{t=1}^{T-1} S_{Gt}$ | $E_G = \frac{1}{T} \sum_{t=1, T} E_{Gt}$ |
| Non-Global | $A_N = \frac{1}{T} \sum_{t=1}^T A_{Nt}$ | $D_N = \frac{1}{T} \sum_{t=1}^{T-1} D_{Nt}$ | $S_N = \frac{1}{T} \sum_{t=1}^{T-1} S_{Nt}$ | $E_N = \frac{1}{T} \sum_{t=1, T} E_{Nt}$ |

Elements of the DSE decomposition can be visualized as time series (see Table 2) or as images. For example, just as a variance image with voxels 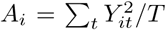 is useful, we find that a *D*-var image, 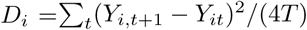 and a *S*-var image, 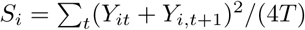 offer more informative views of the noise structure.

### 2.3. DSE ANOVA Table & Reference Values

We see the arrangement of DSE values in Table 1 as a variant of an Analysis of Variance (ANOVA) table that summarizes contributions from fast, slow, end, global and non-global components to the total mean-squares in a 4D dataset. Traditionally ANOVA tables use sum-of-squares to partition variance, but we instead focus on root mean squared (RMS) or mean squared (MS) values to leverage intuition on typical noise standard deviation (or variance) of resting state fMRI data.

We calculate expected values for each of the DSE values for artifact-free data using different null models. In Appendix D we detail the most arbitrary version of this model, based only on time-constant spatial covariance, Σ*^S^*. Another model is based on time-space-separable correlation; this noise model assumes data with arbitrary spatial covariance Σ*^S^* but a common temporal autocorrelation for all voxels with a constant lag-1 autocorrelation *ρ.* While this is a less restrictive time series model than AR(1), in practice temporal autocorrelation varies widely over space, and we stress we only consider this as a working model to gain intuition on the DSE ANOVA table. (Our null model for DVARS p-values, below, does not assume time-space separability). We also consider the idealized model of “perfect” data with completely independent and identically distributed (IID) 4D data.

**Table 2:** Expressions that make up the time series visualization of the DSE variance decomposition. *A*-var is to the total variance at time point *t*, *D*-var, S-var and E-var correspond to the fast, slow and edge variance terms. Global and non-global variance components sum to the total components. All of these terms, given as mean squared quantities, are best reported and plotted in root mean squared (RMS) units (see Appendix B for more on plotting global variance).

| Name | Notation | Value | Range | X-axis Loc. |
| --- | --- | --- | --- | --- |
| A-var | $A_t$ | $\frac{1}{I} \sum_{i=1}^I Y_{it}^2$ | $t = 1, \dots, T$ | $t$ |
| *D*-var | $D_t$ | $\frac{1}{I} \sum_{i=1}^I (Y_{it} - Y_{i,t+1})^2/4$ | $t = 1, \dots, T-1$ | $t + \frac{1}{2}$ |
| *S*-var | $S_t$ | $\frac{1}{I} \sum_{i=1}^I (Y_{it} + Y_{i,t+1})^2/4$ | $t = 1, \dots, T-1$ | $t + \frac{1}{2}$ |
| *E*-var | $E_t$ | $\frac{1}{I} \sum_{i=1}^I Y_{it}^2/2$ | $t = 1, T$ | $t$ |
| Global A-var | $A_{Gt}$ | $\bar{Y}_t^2$ | $t = 1, \dots, T$ | $t$ |
| Global *D*-var | $D_{Gt}$ | $(\bar{Y}_t - \bar{Y}_{t+1})^2/4$ | $t = 1, \dots, T-1$ | $t + \frac{1}{2}$ |
| Global *S*-var | $S_{Gt}$ | $(\bar{Y}_t + \bar{Y}_{t+1})^2/4$ | $t = 1, \dots, T-1$ | $t + \frac{1}{2}$ |
| Global *E*-var | $E_{Gt}$ | $\bar{Y}_t^2/2$ | $t = 1, T$ | $t$ |
| Non-Global A-var | $A_{Nt}$ | $\frac{1}{I} \sum_i (Y_{it} - \bar{Y}_t)^2$ | $t = 1, \dots, T$ | $t$ |
| Non-Global *D*-var | $D_{Nt}$ | $\frac{1}{I} \sum_i (Y_{it} - Y_{i,t+1} - (\bar{Y}_t - \bar{Y}_{t+1}))^2/4$ | $t = 1, \dots, T-1$ | $t + \frac{1}{2}$ |
| Non-Global *S*-var | $S_{Nt}$ | $\frac{1}{I} \sum_i (Y_{it} + Y_{i,t+1} - (\bar{Y}_t + \bar{Y}_{t+1}))^2/4$ | $t = 1, \dots, T-1$ | $t + \frac{1}{2}$ |
| Non-Global *E*-var | $E_{Nt}$ | $\frac{1}{I} \sum_i (Y_{it} - \bar{Y}_t)^2/2$ | $t = 1, T$ | $t$ |

Table 3 shows three sets of reference values for the DSE ANOVA table ^1^. The first pair of rows shows the expected value of the MS for each component for the separable model. This shows that all DSE components scale with the average voxel-wise variance 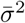, and as temporal autocorrelation *ρ* increases *D*-var shrinks and *S*-var grows. The global components are seen to depend on 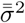, the average of the *I*^2^ elements of Σ*^S^*. This indicates, intuitively, that the greater the spatial structure in the data the more variance that is explained by the global.

The next pair of rows in Table 3 show the expected MS values normalized to the expected *A-*var term. The *A*-var-normalized *D*-var and *S*-var diverge from 1/2 exactly depending on *ρ,* specifically *S-D = ρ*(*T-* 1)/*T*. The global terms here depend on the ratio of average spatial covariance and average variance, 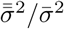.

**Table 3:**
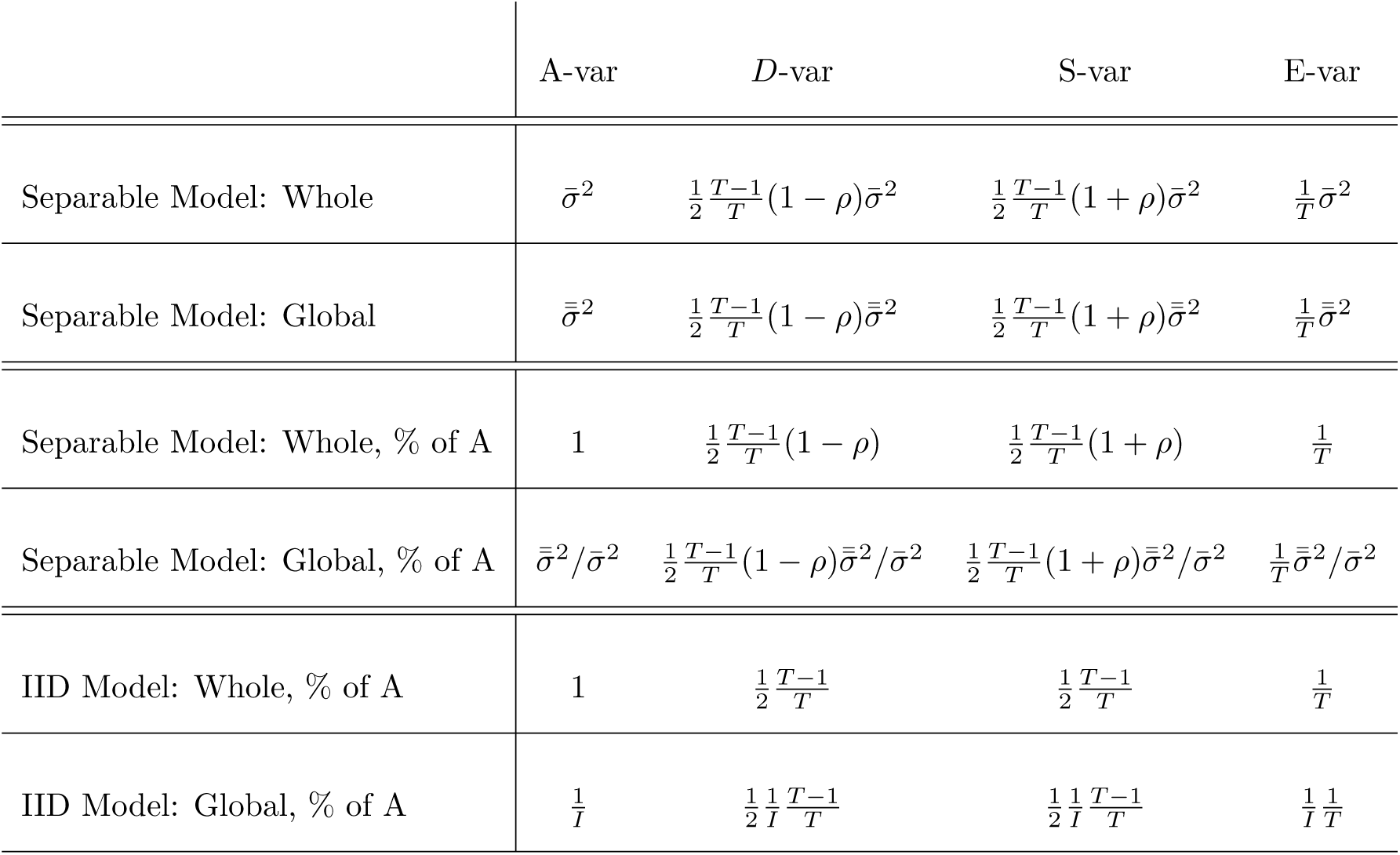
Expected values of the DSE ANOVA table under different nominal models. First two rows show expected mean squared (MS) values under the separable noise model, for whole and global variance. Third and fourth rows show expected MS normalized to the total variance *A*-var for the separable model. Final two rows show the expected normalized MS under a naive, default model of independent and identically distributed (IID) data in time and space. 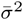 is the average of the *I* voxel-wise variances, *ρ* is the common lag-1 autocorrelation, and 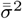 is the average of the *I*^2^ elements of the voxels-by-voxels spatial covariance matrix. This shows that *D*-var and *S*-var are equal under independence but, when normalized, differ by about *ρ*; this is a general result that doesn’t depend on the separable noise model used here (see Appendix D.8).

The final pair of rows shows expected values under the most restrictive case of IID noise. Here *D*-var and S-var are exactly equal, about 1/2, and we see that the global variance explained should be tiny, 1*/I.* This suggests that normalized global variance relative to the nominal IID value, i.e. (*A*_*G*_/*A*) /(1/*I*), an estimate of 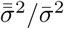, can be used as a unitless index of the strength of spatial structure in the data. (This particular result doesn’t depend on the separable model; see Appendix D).

The handy result on the *S – D* approximating *ρ* generalizes beyond the time-space-separable model: For an arbitrary model, both *S – D* and *S*_*t*_ – *D*_*t*_ normalized to *A* estimate a weighted average of the lag-1 temporal autocorrelations (see Appendix D.8). Hence, the convergence of *D*-var and *S*-var we observe as data is cleaned up has the specific interpretation of reduction in the average lag-1 autocorrelation.

These reference models provide a means to provide DSE values in three useful forms. For each *A*-var, *D*-var, *S*-var and *E*-var term we present:

1. RMS, the square root of the mean squared variance quantity,
2. %A-var, a variance as a percentage of total mean-square *A,* and
3. Relative IID, *A*-var-normalized values in ratio to nominal IID values.

For example, for *A*-var we have (1) RMS is 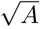, (2) %*A*-var is 100% and (3) relative IID is 1.0. For *D*-var, (1) RMS is 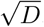, (2) %A-var is *D*/*A ×* 100 and (3) relative IID is

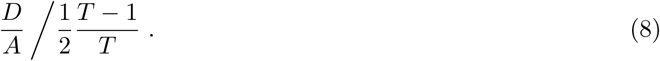

For *D*_*G*_-var, (2) RMS is 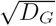, (2) %A-var is *D*_*G*_/*A ×* 100 and (3) relative IID is

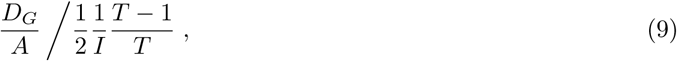

noting that we normalize to *A* and not *A_G_.*

We note that the fast and slow components can be defined as responses of linear time-invariant filters. The slow component corresponds to an integrator filter with power transfer function 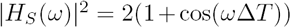 and the fast component corresponds to a differentiator filter with power transfer function 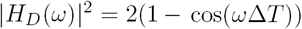, where *ω* is angular frequency and ∆*T* is the repetition time (TR). In other words, in time domain, *S*_*t*_ can be interpreted as average of convolved BOLD signals with a rectangular window of [1 1] and *D*_*t*_ with a [1 −1] window.

### 2.4. Inference for DVARS

We seek a significance test for the null hypothesis

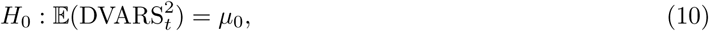

where *µ*_0_ is the mean under artifact-free conditions. Note this is equivalent to a null of homogeneity for DVARS_*t*_ or *D_t_.* If we further assume that the null data are normally distributed, we can create a *χ*^2^ test statistic

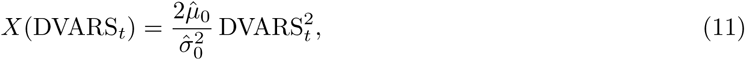

approximately following a 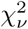 distribution with 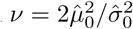 degrees of freedom, where 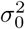 is the null variance (see Appendix E).

What remains is finding estimates of 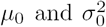. The null mean of DVARS_*t*_ is the average differenced data variance,

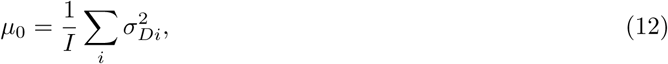

where 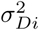 is the variance of the differenced time series at voxel *i.* To avoid sensitivity to outliers, we robustly estimate each 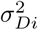 via the interquartile range (IQR) of the differenced data,

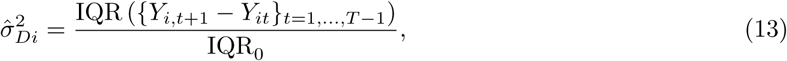

where 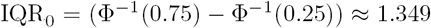 is the IQR of a standard normal, and Φ^−1^(·) is the inverse cumulative distribution function of the standard normal. Below we evaluate alternate estimates of *µ*_0_, including the median of 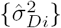 and directly as the median of 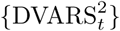.

The variance of 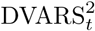 unfortunately depends on the full spatial covariance, and thus we’re left to robustly estimating sample variance of 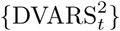 directly. We consider several estimates based on IQR and evaluate each with simulations below. Since the IQR-to-standard deviation ratio depends on a normality assumption, and we consider various power transformations before IQR-based variance estimation (see Appendix F). We also consider a “half IQR” estimate of variance

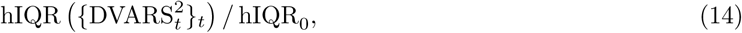

where hIQR is the difference between the median and first quartile, and hIQR_0_ = IQR_0_ /2. This provides additional robustness against contamination of the variance estimate from upward spikes.

Finally, the *X*(DVARS_*t*_) values can be converted to p-values *P*(DVARS_*t*_) with reference to a 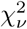 distribution, and subsequently converted into equivalent Z scores,

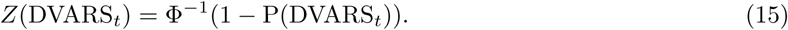

Note that for extremely large values of DVARS_*t*_ numerical underflow will result in p-values of zero; in such cases an approximate Z score can be obtained directly as 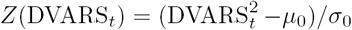.

Under complete spatial independence the degrees of freedom will equal the number of voxels *I,* and so *ν* can be thought of an effective number of spatial elements; large scale structure will decrease *ν* while larger *ν* should be found with cleaner data. Though we caution that estimates of *ν* will be very sensitive to the particular estimators used for 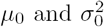.

### 2.5. Standardized DVARS

For intra-cohort investigation of corruptions, we propose that our *D*-var time series, 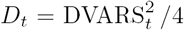, is a more interpretable variant of DVARS, as it represents a particular “fast” portion of noise variance, and when added to “slow” mean-square, *S*_*t*_, gives the total mean-square of the 4D data 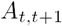. However, these components are not suitable for inter-cohort comparisons, as the variance characteristics may vary with acquisition or scanner differences. In this section we propose a set of transformations which makes the inter-cohort comparison of the DSE components (including DVARS) possible.

First consider the percent *D*-var variance explained at a single time point. Eqn. (5) could be used to find, in sums-of-squares units, the percent variance attributable to *D*-var at *t, t +* 1:

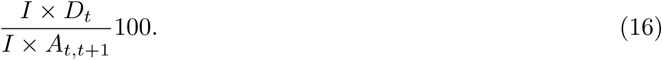

However, problem scans can inflate *A*_*t*_ and could mask spikes. Hence we instead propose to replace *A*_*t*_,_*t*+1_ with its average *A* and compute percent *D*-var at time *t* as

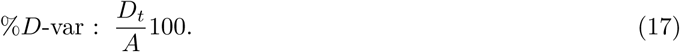

This has the merit of being interpretable across datasets, regardless of total variance. This is just percent normalization to *A* as discussed above.

While %*D*-var can be more interpretable than unnormalized *D*-var, its overall mean is still influenced by the temporal autocorrelation. For example, if %*D*-var is overall around 30% and at one point there is a spike up to 50%, what is interesting is the 20 percentage point change, not 30% or 50% individually. Hence another useful alternative is change in percent *D*-var from baseline

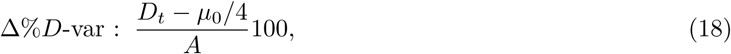

interpretable as the excess fast variability as a percentage of average variance. Later in Section 4.2.1, we show how ∆%*D*-var is used as measure of “practical significance” to complement DVARS p-values.

We previously have proposed scaling DVARS relative to its null mean (Nichols, 2013),

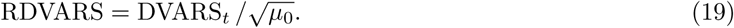

(While we had called this “Standardized DVARS”, a better label is “Relative DVARS.”) This gives a positive quantity that is near 1 for good scans and substantially larger than one for bad ones. However, there is no special interpretation “how large” as the units (multiples of 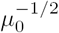) are arbitrary; as noted above, DVARS falls with increased temporal correlation, making the comparison of these values between datasets difficult.

Finally the Z-score Z(DVARS_*t*_) or − log_10_ P(DVARS_*t*_) may be useful summaries of evidence for anomalies.

## 3. Methods

### 3.1. Simulations

To validate our null distribution and p-values for DVARS we simulate 4D data as completely independent 4D normally distributed noise

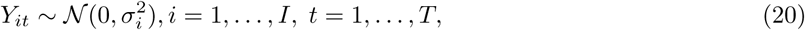

for *σ*_*i*_ drawn uniformly between *σ*_min_ and *σ*_max_ for each *i, I =* 90, 000.

We manipulate two aspects in our simulations, time series length and heterogeneity of variance over voxels. We consider *T* of 100, 200, 600 and 1200 data-points, reflecting typical lengths as well as those in the Human Connectome Project. We use three variance scenarios, homogeneous with *σ*_min_ = *σ*_max_ = 200, low heterogeneity *σ*_min_ = 200 and *σ*_max_ = 250, and high heterogeneity *σ*_min_ = 200 and *σ*_max_ = 500.

We consider four estimates of *μ*_0_ First is the very non-robust sample mean of 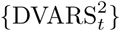, denoted 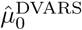, considered only for comparative purposes. Next we compute the mean 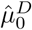 and median 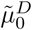 of 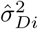 (Eqn. (13)), the robust IQR-based estimates of differenced data variance at each voxel. Finally we also consider the empirical median of 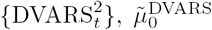. For 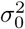, all estimates were based directly on 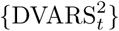; for comparative purposes we considered the (non-robust) sample variance of 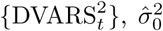, and IQR-based and hIQR-based estimates of variance with power transformations *d* of 1, 1/2, 1/3 and 1/4, denoted generically 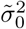; note *d* = 3 is theoretically optimal for 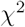 (see Appendix F).

For p-value evaluations, we only evaluate the most promising null moment estimators 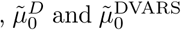 for 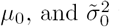 with hIQR, *d =* 1 and hIQR, *d =* 3. We measure the bias our estimators in percentage terms, as 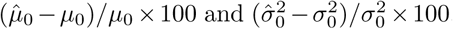, where the true value are 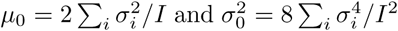 (as per Appendix E when Σ^*s*^ = *I*). For each method we obtain P-values and create log P-P plots (probability-probability plots) and histograms of equivalent Z-scores.

Similar simulation settings are used to evaluate the power of the DVARS hypothesis test, except we consider 4 different autocorrelation levels *ρ =* {0,0.2, 0.4,0.6}. This range is chosen to reflect observed estimates of lag-1 autocorrelation coefficients in the HCP cohort. Inferences are assessed in terms of sensitivity and specificity.

All simulations use 1,000 realisations.

### 3.2. Analysis of Functional Connectivity

We evaluate the impact of the DVARS test as a tool for “scrubbing” (scan deletion) on functional connectivity (FC) measued with Pearson’s correlation coefficient. We consider FC between all possible pairs of Region of Interests (ROI) in each subject for a given ROI atlas. The mean time series of each ROI is obtained by averaging all the time series within a ROI. To parcellate the brain, we use two data-driven atlases; Power Atlas (Power et al., 2011) which is constructed of 264 non-neighboring cortical and sub-cortical ROIs and each ROIs has 81 voxels (is case of 2mm isotropic volumes) and Gordon Atlas (Gordon et al., 2014) which is constructed of 333 cortical regions of interests with different sizes.

We use two popular methods to evaluate the effect of the DVARS inference on functional connectivity. First, we use the QC-FC analysis which begins by creating per-edge, intersubject scores, the correlation of the number of removed volumes and FC; these scores are plotted against the inter-ROI distance (in mm). We then use LOESS smoothing method (with span window of %1) to summarize the association for each method. For further details about QC-FC method, see Power et al. (2014a); Ciric et al. (2016); Burgess et al. (2016). We use QC-FC to compare our DVARS hypothesis test to four other scan scrubbing methods. From Power et al. (2012) we use two FD thresholds, lenient (0.2mm) and conservative (0.5mm), and a DVARS threshold of 5. From FSL’s fsl motion outliers tool (Jenkinson et al., 2012), we use a DVARS threshold corresponding to box-plot right-outliers, 1.5 IQRs above the 75%ile. Note that the first three approaches used a fixed threshold, while the FSL approach gives a run-specific threshold.

The objective of this FC analysis is to investigate whether DVARS inference test performs as well as the available thresholding methods (such as arbitrary thresholding of FD (Power et al., 2012) and DVARS (Burgess et al., 2016)) and if so, whether it delivers the optimal results while sacrificing the fewest temporal degree of freedom as possible. Therefore, we only present the results for the Minimally pre-processed data sets.

### 3.3. Real Data

We use two publicly available data-sets to demonstrate the results of methods proposed in this paper on real-data. First, we use 100 subjects from ”100 Unrelated” package in the Human Connectome Project (HCP,S1200 release). We chose this dataset due to the high quality and long sessions of the data (Smith et al., 2013; Glasser et al., 2013). Second, we used first 25 healthy subjects from the New York University (NYU) cohort of the Autism Brain Imaging Data Exchange (ABIDE) consortium via Preprocessed Connectome Project (PCP) (Craddock et al., 2013). We selected this cohort for its high signal-to-noise ratio and the more typical (shorter) time series length (Di Martino et al., 2014).

#### 3.3.1. Human Connectome Project (HCP)

For full details see (Van Essen et al., 2013; Glasser et al., 2013); in brief, 15 minute eyes-open resting acquisitions were taken on a Siemens customized Connectome 3T scanner with a gradient-echo EPI sequence, TR=720ms, TE=33.1 ms, flip angle=52° and 2 mm^3^ isotropic voxels. For each subject, we used the first session, left to right phase encoding direction (See Table S1 for full details of subjects). We considered each subject’s data in three states of pre-processing: unprocessed, minimally pre-processed and ICA-FIXed processed. Unprocessed refers to the raw data as acquired from the machine without any pre-processing step performed, useful as a reference to see how the DSE components change with preprocessing steps. Minimally pre-processed (MPP) data have undergone a range of conventional pre-processing steps such as correction of gradient-nonlinearity-induced distortion, realignment aiming to correct the head movements, registration of the scans to the structural (T1w) images, modest (2000s) high pass filtering and finally transformation of the images to the MNI standard space.

Finally, after regressing out the 24-motion parameters, an ICA-based clean up algorithm called ICA-FIX (Salimi-Khorshidi et al., 2014) is applied, where artifactual ICA components, such as movement, physiological noises of the heart beat and respiration, are regressed out non-aggressively. Due to extent of the FIX denoising and an ongoing debate regarding the nature of the global signal, we did not consider global signal regression with the HCP data. From now on, we call this stage ‘fully pre-processed (FPP)’ to be consistent with the ABIDE-NYU cohort we describe in the following.

#### 3.3.2. Autism Brain Imaging Data Exchange (ABIDE)

We use use 20 healthy subjects of New York University (NYU) data-set. For full details visit Pre-processed Connectome Project website http://preprocessed-connectomes-project.org/; in brief, 6 minute eyes-closed resting acquisitions were taken on an Allegra 3T scanner with a gradient echo EPI sequence, TR=2000ms, TE=15ms, flip angle=90°, and 3 mm isotropic voxels (See Table S2 for full details of subjects). In this study, each subject was analyzed using Configurable Pipeline for the Analysis of Connectomes (C-PAC) pipeline, in three stages; unprocessed, minimally pre-processed and fully pre-processed. The unprocessed data are raw except for brain extraction with FSL’s BET. Minimally pre-processed data were only corrected for slice timing, motion by realignment and then the data were transformed into a template with 3 mm^3^ isotropic voxels. Fully pre-processed data additionally had residualisation with respect to 24-motion-parameters, signals from white matter (WM) and cerebrospinal fluid (CSF), and linear and quadratic low-frequency drifts. Conventionally this pipeline deletes the first three volumes to account for T1 equilibration effects, but we examine the impact of omitting this step for the raw data.

Further, we also use all healthy subject of ABIDE (530 subjects) to show how DSE decomposition can be used to compare the data-sets, cohorts and pipelines.

## 4. Results

### 4.1. Simulations

Figure 2 shows the percentage bias for the null expected value *μ*_0_ (left panel) and variance 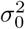 (right panel) for different levels of variance heterogeneity and time series length.

**Figure 2:**
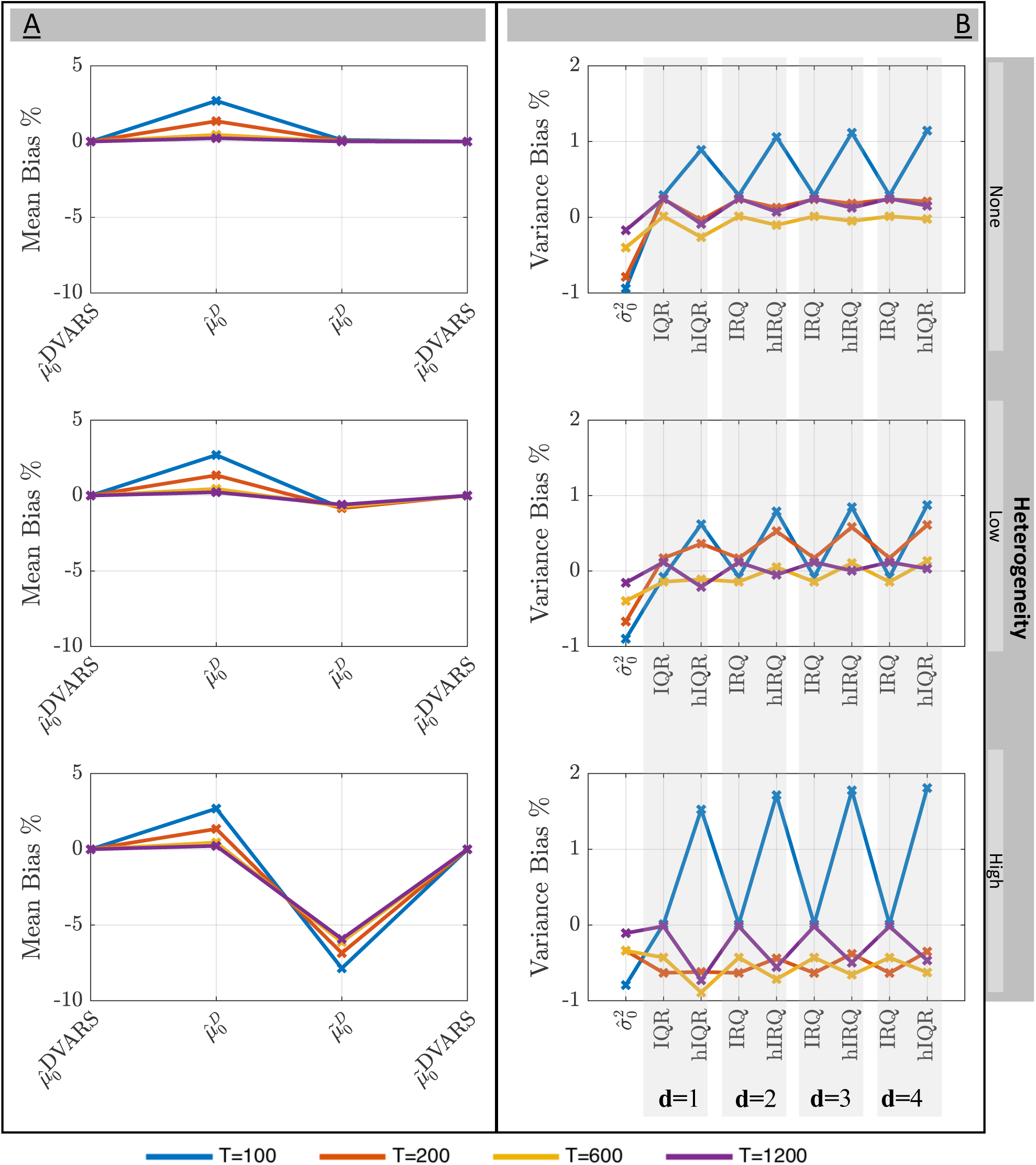
Simulation results for estimation of mean and variance of DVARS^2^ under the null of temporal homogeneity. The mean *μ*_0_ (left) and variance 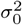 (right) are shown for no, low and high spatial heterogeneity of variance (rows). All estimators improve with time series length *T* and most degrade with increased spatial heterogeneity. For the mean, both the sample mean 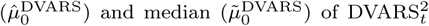 perform well, as does voxel-wise median of difference data variance 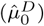 for sufficient *T,* though 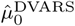 of course lacks robustness. For *T* ≥ 200, all variance estimators have less than 1% bias.

The direct estimates of the *μ*_0_ based on the 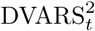 time series perform best on this clean, artifactfree data, while *μ*_0_ estimated on variance of the differenced data 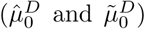 degrades with increasing heterogeneity. The estimates of variance have relatively less bias but it is difficult to identify one particular best method, save for IQR often (but not always) having less bias than hIQR, and lower *d* generally associated with less bias.

On balance, given the generally equivocal results and concerns about robustness, for further consideration we focus on 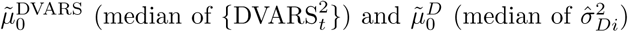 as promising candidates for *μ*_0_, and hIQR with *d =* 1 and hIQR with *d =* 3 for 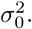.

Figure 3 shows log P-P plots for 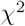 p-values and histograms of approximate Z scores, 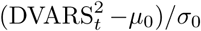; values above the identity in the P-P plot correspond to valid behavior. While all methods have good performance under homogeneous data, 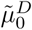 (panels A & C) is not robust to variance heterogeneity and results in inflated significance. In contrast, 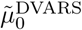 (panels B & D) has good performance over all, for variance estimated with either *d* = 1 or *d* = 3 (top and bottom panels, respectively), and also yields good approximate Z-scores. On the basis of these results, we elected to use 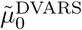 as the only reliable option for the mean, and hIQR, *d* = 3 as a variance estimate, and use these settings going forward.

**Figure 3:**
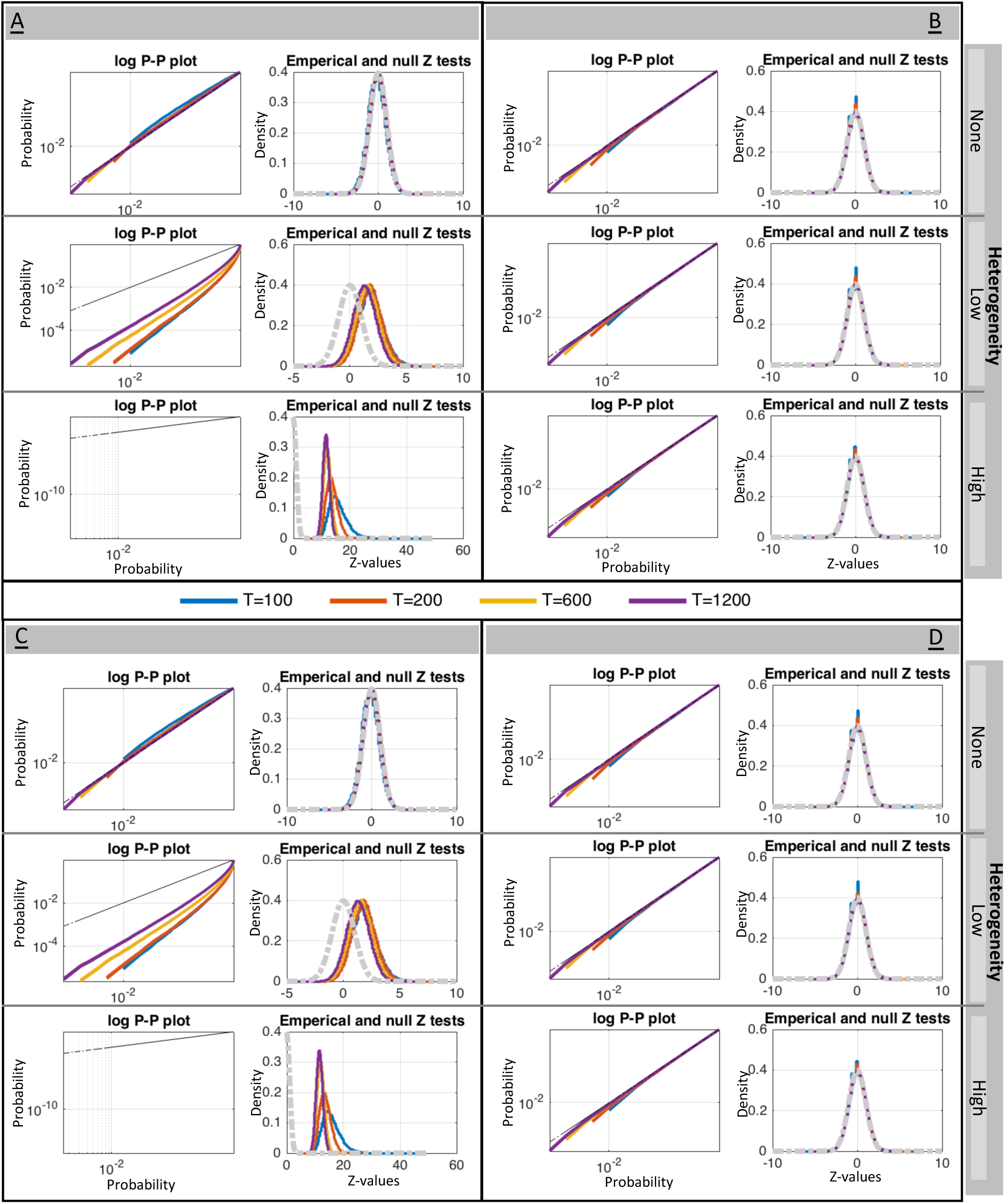
Simulation results for the validity of DVARS p-values for different estimators of *μ*_0_ and 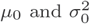. The left two panels (A & C) use 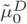, the two right panels (B & D) use 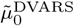; the upper two panels (A & B) use variance based on hIQR with *d =* 1, the lower two panels (C & D) use hIQR with *d =* 3. P-P plots and histograms of Z scores show that only use of 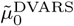 gives reliable inferences, and that the power transformation parameter *d* seems to have little effect.

Figure 4 shows the results of the power simulation. For all sample sizes and autocorrelation parameters, and for the 1% and 10% artifact rates, power was always above 80% and often ≈100%. Increased autocorrelation resulted in improvements in power, while higher artifact rates reduced power. For the 20% artifact rate power was adequate (≈ 80%), but falls to zero for the 30% artifact rate. These results suggest that, at the highest spike rate, the artifacts start to be become indistinguishable from the overall noise (see Fig. S2 for one realization). However, the distribution of DVARS values (Fig. S1) suggest that the constituent null and artifact components are distinguishable even at the highest spike rate, but would require yet more robust methods for estimating the null component than we have employed.

**Figure 4:**
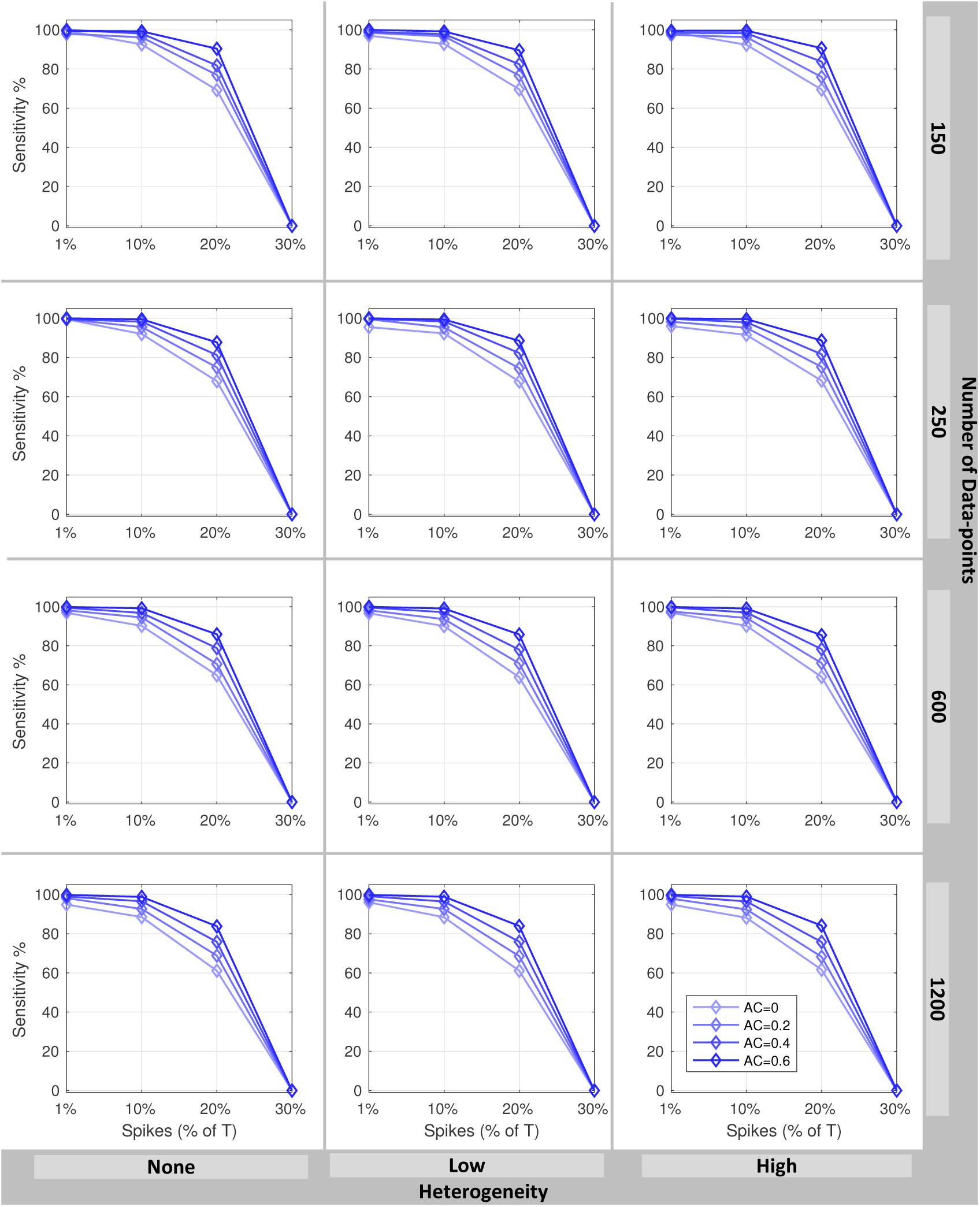
Power of the DVARS hypothesis test to detect artifactual spikes. Plots show sensitivity (% true spikes detected) versus number of true spikes as a percentage of time series length *T,* for varying degrees of temporal autocorrelations (line color). Different *T* (rows) and degree of spatial variance heterogeneity (columns) are considered. These results show hat power increases with autocorrelation but falls with increasing prevalance of spikes; for up to 10% spikes we have excellent power, and for 20% spikes we have satisfactory power (60–90% sensitivity).

### 4.2. Real Data

We first focus on selected results of two HCP subjects, then later summarize results for all HCP and ABIDE subjects.

#### 4.2.1. Temporal Diagnostics: DVARS Inference and Standardized Measures

Figure 5 shows different standardized DVARS measures, as introduced in section 2.5, as well as the other DSE components for subject 118730 of the HCP cohort (See Figs. S4, S5 and S6 for more results.). The first six plots corresponds to the variants listed in Table 4; the bottom two plots show “DSE plots,” plots of *A*_*t*_, *D*_*t*_, *S*_*t*_ and *E*_*t*_ components, upper plot with minimal pre-processing, lower with full pre-processing. The gray and magenta stripes indicate 19 data points identified as having significant DVARS after Bonferroni correction, with magenta indicating time-points that are additionally practically significant by the criterion ∆%*D*-var > 5%. In Figure 5, the largest *D*_t_ occurs at index 7 (i.e. 7th and 8th data points) and has 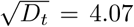, large in terms of being %*D*-var=70.16% of average variance, *Z* = 36.33 indicating extreme evidence for a spike, and having ∆%*D*-var = 41.20% more sum-of-squares variability than expected. The least significant *D*_*t*_ occurs at index 726, with 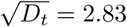; while its *Z* = 4.36 is not a small Z-score, with just ∆%*D*-var=4.95% excess variation, it is a relatively modest disturbance. In contrast, we find that the values of original DVARS or relative DVARS do not offer a meaningful interpretation. Table S3 shows values for all significant scans.

**Figure 5:**
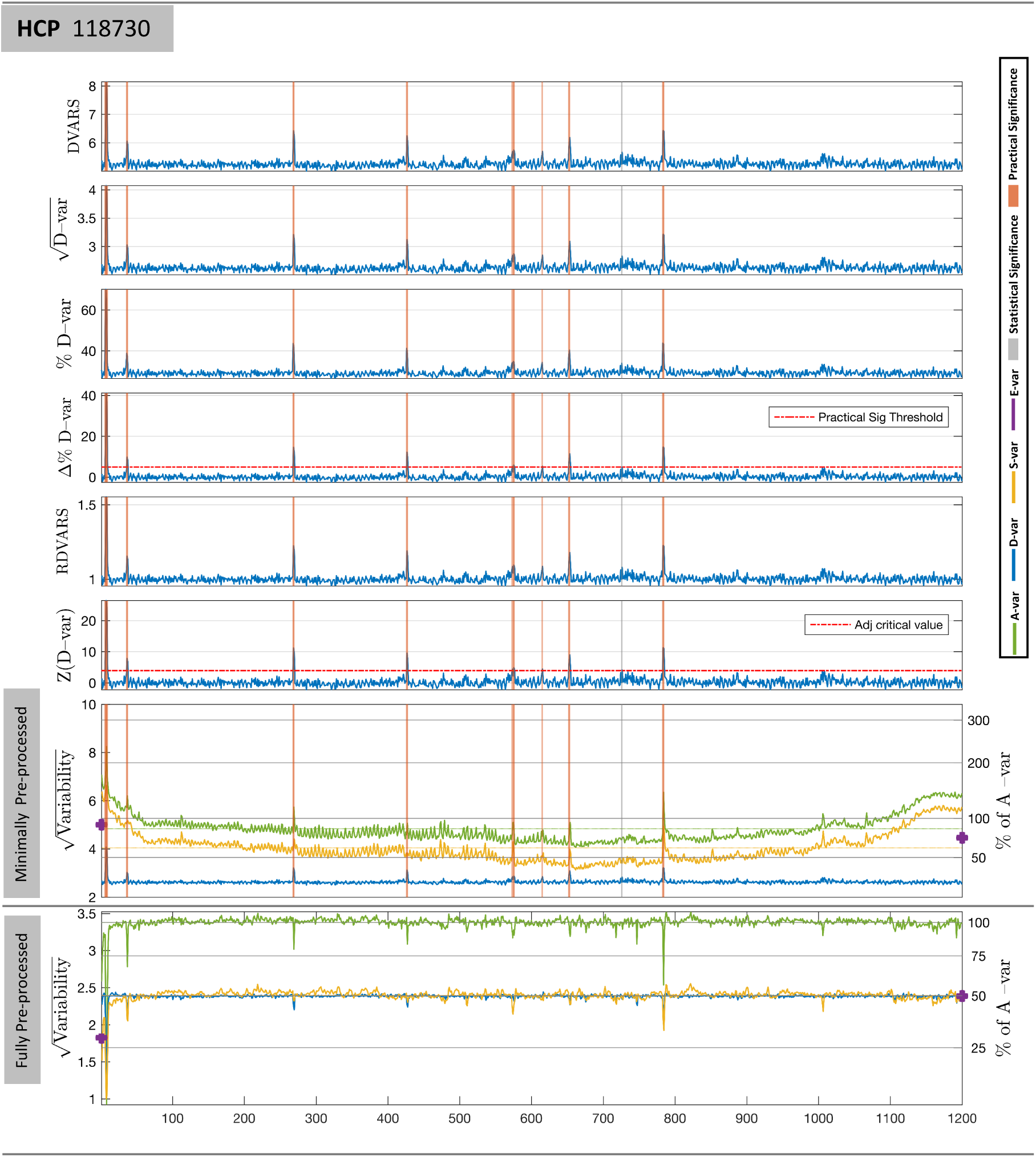
Comparison of different variants of DVARS-related measures on HCP 115320. The first six plots are variants of DVARS listed in Table 4; A%*D*-var is marked with a practical significance threshold of 5%, and Z(DVARS) with the onesided level 5% Bonferroni significance threshold for 1200 scans. Vertical grey stripes mark scans that only attain statistical significance, while orange stripes mark those with both statistical and practical significance. The bottom two plots show the 4 DSE components, total *A*_*t*_ (green), fast *D*_*t*_ (blue), *S*_*t*_ slow (yellow), and edge *E*_*t*_ (purple), for minimally preprocessed (upper) and fully preprocessed (lower) data. For minimally preprocessed data *D*-var is about 25% of *A*-var (see right axis), far below *S*-var. For fully preprocessed data *D*-var and *S*-var converge to 50%*A*-var.

**Table 4:** Form and interpretation of various DVARS variants, expressed as functions of original DVARSt. Here {*Y*_*it*_} are the 4D data, *A* is the overall mean square variance, *µ*_0_ is the expected 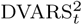 under a null model, P(DVARS_*t*_) is the p-value for DVARS^2^, and Φ^−1^ is the inverse cumulative distribution function of a normal.

| Name | Expression | Interpretation |
| --- | --- | --- |
| DVARs | $D\text{VAR}_t = \sqrt{\sum_i (Y_{it} - Y_{i,t+1})^2 / I}$ | Standard deviation of difference image |
| $\sqrt{D}$-var | $D\text{VAR}_t / 2$ | Fast component of noise, as standard deviation |
| %$D$-var | $D\text{VAR}_t^2 / (4A) \times 100$ | Fast noise, as % of average noise variance |
| $\Delta\%$ $D$-var | $(D\text{VAR}_t^2 - \mu_0) / (4A) \times 100$ | Excess fast noise, as % of average noise variance |
| Rel. DVARs | $D\text{VAR}_t / \sqrt{\mu_0}$ | DVARs as a multiple of null mean |
| $Z(D\text{-var})$ | $\Phi^{-1}(1 - P(D\text{VAR}_t))$ | DVARs p-value as Z-score |

The bottom panel of Figure 5 shows the DSE plot for fully pre-processed data. This data now exhibits the idealized behavior of IID data, with *D*-var and *S*-var components converging at 50% of average variance (see right-hand y-axis). However, interestingly, the change is not similar for all DSE components. Note how 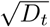 is around 2.6 before clean up, and 2.5 after clean up, while 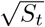 falls dramatically with cleaning, indicating that nuisance variance removed was largely of a “slow” variety. Also observe that clean up results in drops in total *A*_*t*_ variance where artifacts were observed, indicating variance removed by FIX.

Finally, Table 5 explores the use of the estimated 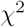 degrees of freedom *v* as an index of spatial effective degrees of freedom. Raw data, exhibiting substantial spatial structure, has *v =* 287, which increases to *v =* 11,086 for fully preprocessed data, still only about 5% of the actual number of voxels.

**Table 5:** Spatial effective degrees of freedom (EDF) for HCP subject 115320. As more spatial structure is removed with preprocessing, spatial EDF rises, but never to more than 5% of the actual number of voxels.

|  | Voxels | Spatial EDF | Spatial EDF / Voxels |
| --- | --- | --- | --- |
| Raw | 162,768 | 287 | 0.176% |
| MPP | 224,998 | 1,660 | 0.738% |
| FPP | 224,998 | 11,086 | 4.928% |

#### 4.2.2. Effect of DVARS Inference Testing on Functional Connectivity

FC evaluations based on 55,278 unique edges from the Gordon atlas are shown in Figure 6 (see Fig. S9 for Power Atlas results). Panel A shows the QC-FC analysis of five thresholding methods, compared to unscrubbed QC-FC. The results from the DVARS test appear comparable to the other methods, but Panel B of Figure 6 show that the DVARS test removes many fewer scans on average, preserving temporal degrees of freedom. A related evaluation, comparing DVARS hypothesis test scrubbing to random scrubbing, finds that FC is significant impacted by the DVARS scrubbing (Fig. S10 and S11).

**Figure 6:**
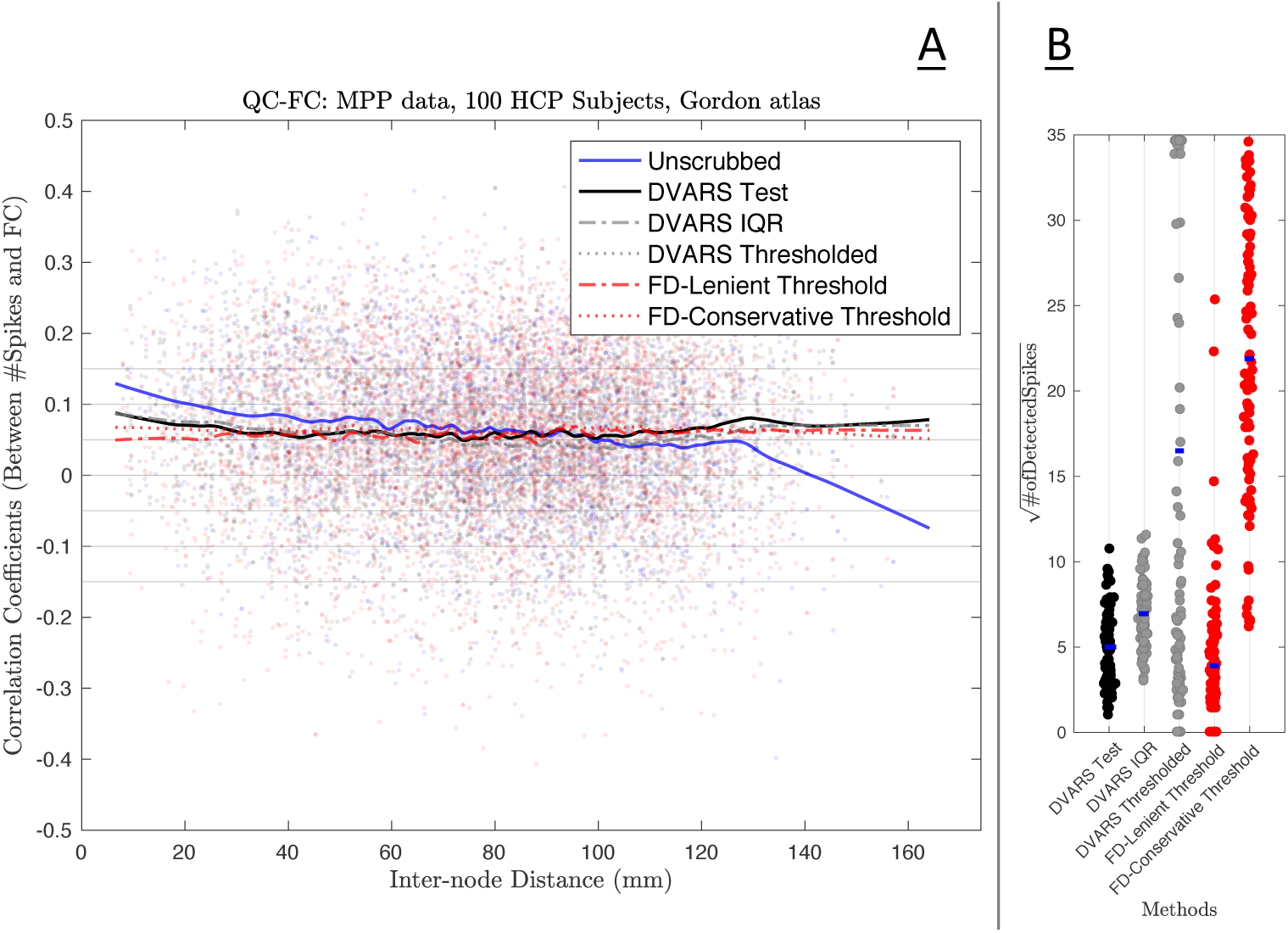
Impact of scrubbing on functional connectivity of 100 HCP subjects’ MPP data, comparing the DVARS test to four other existing methods. Panel A shows the QC-FC analysis for five different thresholding methods (see body text for details); shown are DVARS test, FD thresholding (FD-Lenient & FD-Conservative), arbitrary DVARS threshold, and DVARS boxplot outlier threshold (DVARS IQR). Panel B shows the loss of temporal degree of freedom for each method (i.e. number of scans scrubbed), one dot per subject and dot color following line colors in Panel A. These result show that, in terms of FC, all the methods are largely equivalent, but the DVARS test is best at preserving degrees of freedom.

We note that the sole purpose of preceeding QC-FC analysis is to ensure that the DVARS inference test outperforms other arbitrary thresholds available in literature, and therefore we do not show the similar results for FPP data.

#### 4.2.3. Temporal Diagnostics: Before and After Clean-up

Figures 7 and 8 shows the minimally and fully pre-processed DSE decompositions, respectively, of HCP subject 115320.

**Figure 7:**
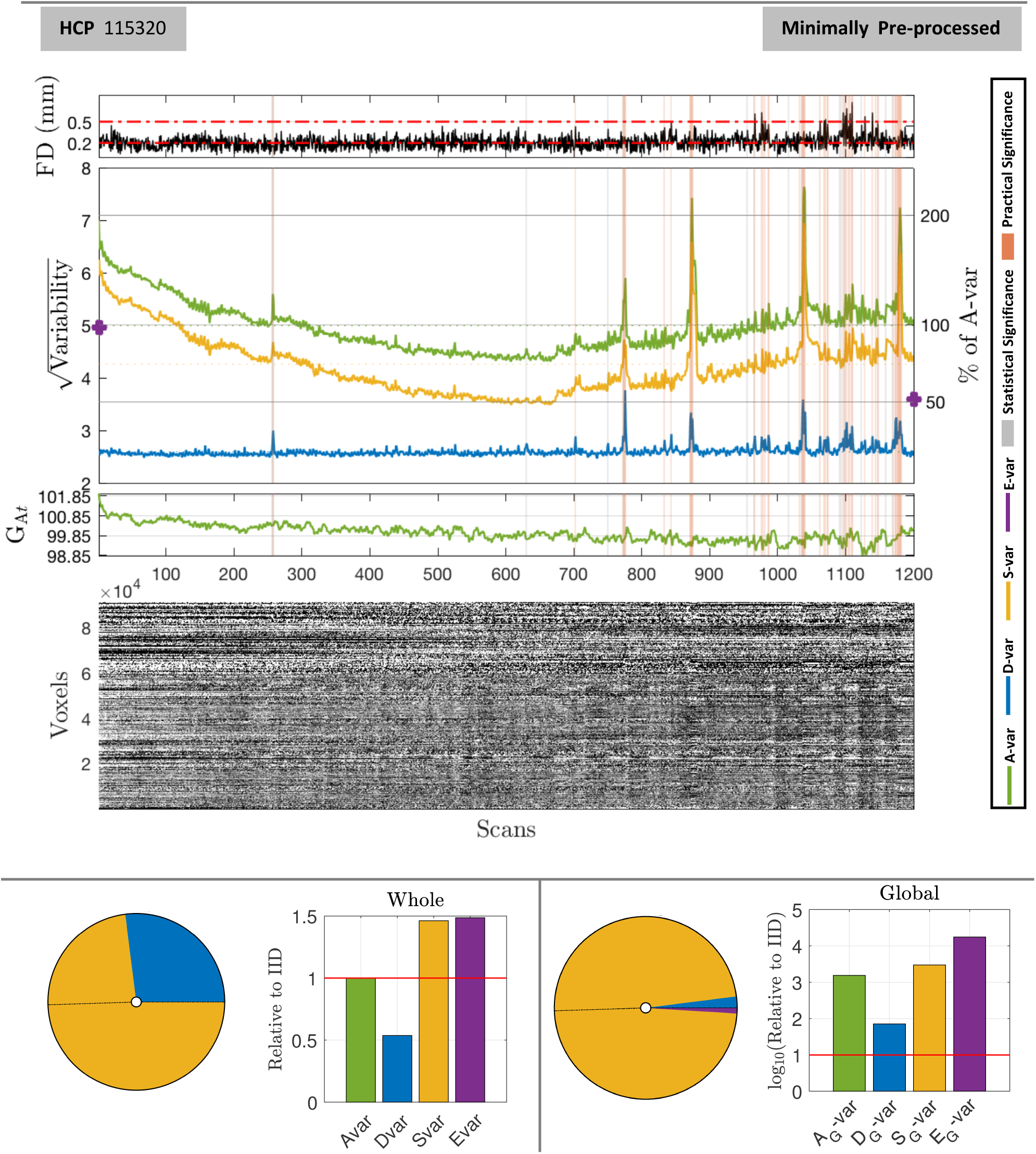
DSE and DVARS inference for HCP 115320 minimally pre-processed data. The upper panel shows four plots, framewise displacement (FD), the DSE plot, the global variance signal G At, and an image of all brainordinate elements. FD plots show the conventional 0.2mm and 0.5mm, strict and lenient thresholds, respectively. All time series plots have DVARS test significant scans marked, gray if only statistically significant (5% Bonferroni), in orange if also practically significant (∆%*D*-var>5%). The bottom panel summaries the DSE ANOVA table, showing pie chart of the 4 SS components and a bar chart relative to IID data, for whole (left) and global (right) components. Many scans are marked as significant, reflecting disturbances in the latter half of the acquisition.

**Figure 8:**
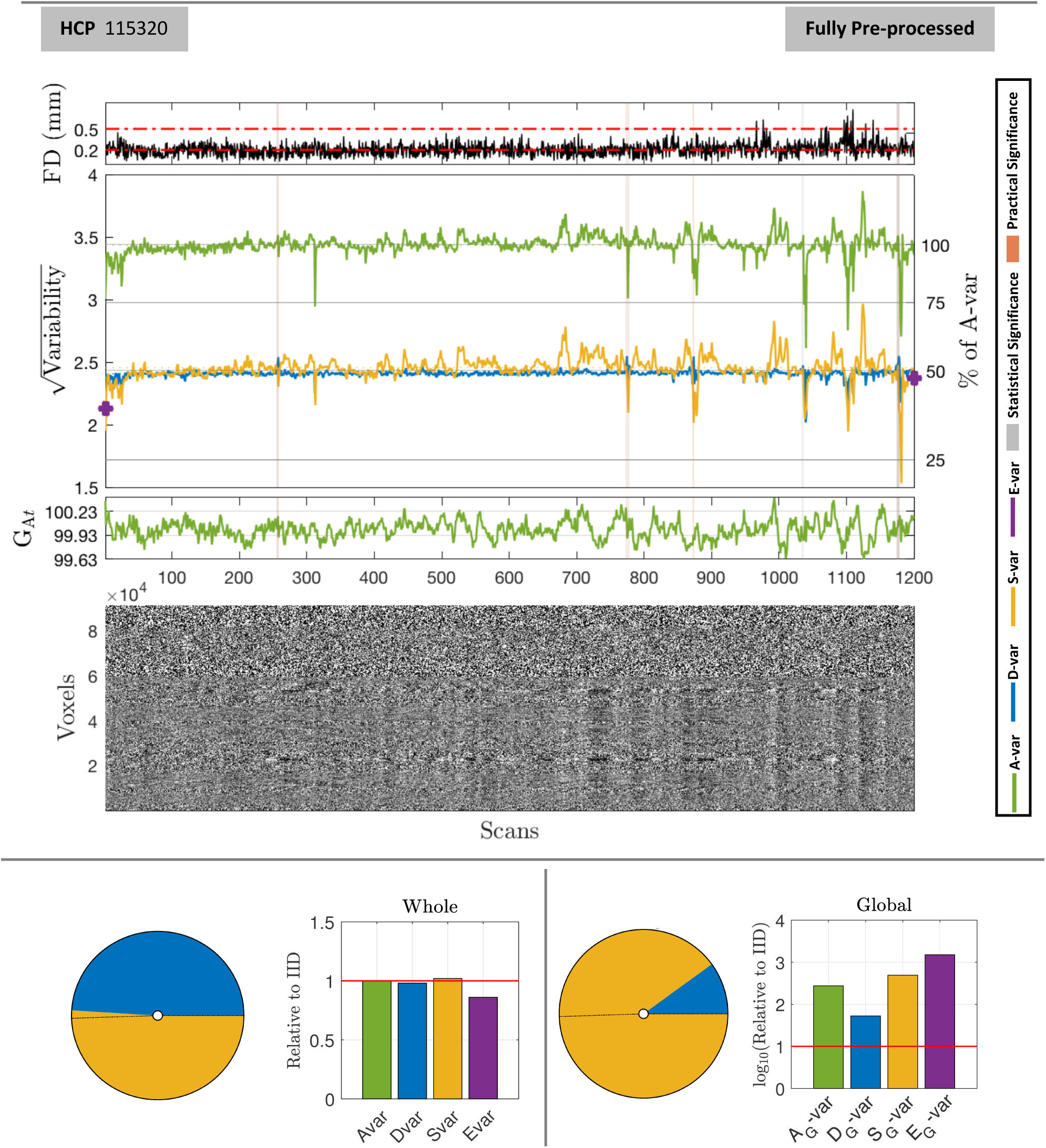
DSE and DVARS inference for HCP 115320 fully pre-processed. Layout as in Figure 7. Cleaning has brought *S*_*t*_ slow variability into line with *D*_*t*_ fast variability, each explaining about 50% of total sum-of-squares. While some scans are still flagged as significant, %*D*-var (*D* as a % of *A*-var, right y axis) never rises above about 55%, indicating ∆%*D*-vars of 5% or less lack of practical significance.

Figure 7, upper panel, shows that if the strict FD threshold, 0.2mm (Power et al., 2014b), were used 47% of scans would be flagged, while the lenient threshold, 0.5mm (Power et al., 2014b), appears to miss several important events. For example, around scans 775 and 875 there are two surges in 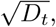 rising to about 60% and 40% average sum-of-squares (excesses of 30% and 10%, respectively, from a baseline of about 30%) while FD remains low. The lower panel’s pie chart shows that *S*-var explains just under 75% of total, and almost all of global sum-of-squares. The Edge component is also 1.5 above its expectation.

In Figure 8, the fully preprocessed data-set shows roughly equal fast and slow components, as reflected in the overlapping *D*_t_ and *S*_t_ sum-of-squares time series (blue and yellow, respectively) and the pie and bar charts for total sum-of-squares. Edge component *E*-var has also dropped to fall in line with IID expectations. However, this convergence is not homogeneous over scans and excursions of *S*-var are still found after scan 650. However, these are much reduced relative to MPP data (no more than 75% of average sum-of-squares, compared to over 150% in Fig. 7).

Note that while significant DVARS are found in the FPP data, they are small in magnitude: Table 6 lists the 10 significant tests, none with ∆%*D*-var greater than 6%. If we used a ∆%*D*-var of 5% we would still mark 4 of these 10 significant; while we might hope for better performance from the FIX method, note the severe problems detected towards the end of the scan (Fig. 7).

**Table 6:** List of all statistically significant *D*_*t*_ fast SS components in the fully pre-processed HCP 115320. Spikes which represent the highest (index 1177) and lowest (index 1035) are marked in bold.

| Scan | Index | DVARS | $\sqrt{D\text{-var}}$ | %$D\text{-var}$ | $\Delta\%D\text{-var}$ | RDVARS | Z($D\text{-var}$) | FD |
| --- | --- | --- | --- | --- | --- | --- | --- | --- |
| 256 & 257 | 256 | 4.982 | 2.4910 | 52.519 | 3.362 | 1.038 | 5.093 | 0.136 |
| 257 & 258 | 257 | 5.077 | 2.538 | 54.553 | 5.397 | 1.058 | 8.175 | 0.172 |
| 774 & 775 | 774 | 5.095 | 2.547 | 54.935 | 5.779 | 1.062 | 8.753 | 0.290 |
| 777 & 778 | 777 | 4.955 | 2.477 | 51.950 | 2.794 | 1.033 | 4.232 | 0.247 |
| 873 & 874 | 873 | 5.089 | 2.544 | 54.805 | 5.649 | 1.061 | 8.556 | 0.255 |
| **1035 & 1036** | **1035** | **4.948** | **2.474** | **51.815** | **2.659** | **1.031** | **4.027** | **0.280** |
| 1175 & 1176 | 1175 | 4.960 | 2.480 | 52.062 | 2.905 | 1.034 | 4.401 | 0.109 |
| 1176 & 1177 | 1176 | 4.953 | 2.476 | 51.926 | 2.769 | 1.032 | 4.195 | 0.104 |
| **1177 & 1178** | **1177** | **5.096** | **2.548** | **54.964** | **5.807** | **1.062** | **8.796** | **0.301** |
| 1178 & 1179 | 1178 | 5.049 | 2.524 | 53.952 | 4.795 | 1.052 | 7.263 | 0.132 |

The smallest significant ∆%*D*-var is 2.66%, which is smaller than the least significant scan detected in the minimally preprocessed data, 3.78%. This indicates the increased sensitivity in our procedure as the background noise in the data is reduced. Note that the majority of the spikes detected in Figure 7 has been removed by ICA-FIX (Fig. 8), however the algorithm has left down-spikes which could be detected via a two-sided version of the test explained in section 2.4.

Temporal diagnostics of before and after clean-up for three other subjects (HCP subject 118730, NYU-ABIDE subjects 51050 and 51050) also reported in Supplementary Materials. See Figure S12 and S13 for HCP subject 118730, Figure S14 and S15 for NYU-ABIDE subject 51050 and Figure S16 and S17 for NYU-ABIDE 51055.

The DSE ANOVA tables for minimally and fully preprocessed (Table 7) gives concise summaries of the data quality. The RMS values provide concrete values that can be used to build intuition for data from a given scanner or protocol. The total noise standard deviation falls from 5.015 to 3.437 with clean-up, but it is notable that the fast component, *D*-var, falls only slightly from 2.598 to 2.406 (in RMS units), while slow variability falls dramatically from about 4.287 to 2.454. This indicates that much of the variance reduction in “cleaning” comes from removal of low frequency drifts and other slowly-varying effects. The magnitude of temporally structured noise is reflected by *S*-var explaining 73% of total sum-of-squares, and after clean-up *S*-var and *D*-var fall into line around 50%. A measure of the spatially structured noise is the global *A*_G_-var that, while small as a percentage, is seen to be about 1,500 that expected with IID before preprocessing, and falling to about 275 relative to IID after preprocessing. That the majority of *A*_G_-var is due to *S*_G_-var indicates that the global signal is generally low frequency in nature.

**Table 7:** DSE ANOVA Tables for HCP 115320. Minimally preprocessed data (top), fully preprocessed (bottom) are readily compared: Overall standard deviation drops from 5.015 to 3.437, while fast noise only reduces modestly from 2.598 to 2.406, indicating preprocessing mostly affects the slow variability. The IID-relative values for *D*, *S* and *E* for the fully preprocessed data are close to 1.0, suggesting successful clean-up in the temporal domain; the global signal, however, still explains about 275× more variability than expected under IID settings, indicating the (inevitable) spatial structure in the cleaned data.

| Minimally Preprocessed Data |  |  |  |
| --- | --- | --- | --- |
| Source | RMS | % of A-var | Relative to IID |
| $A$ - All | 5.015 | 100.000 | 1.000 |
| $D$ - Fast | 2.598 | 26.837 | 0.537 |
| $S$ - Slow | 4.287 | 73.039 | 1.462 |
| $E$ - Edge | 0.176 | 0.124 | 1.486 |
| $A_G$ - All Global | 0.415 | 0.684 | 1539.383 |
| $D_G$ - Fast Global | 0.063 | 0.016 | 71.126 |
| $S_G$ - Slow Global | 0.408 | 0.662 | 2,980.787 |
| $E_G$ - Edge Global | 0.040 | 0.006 | 17,636.960 |
| Fully Preprocessed Data |  |  |  |
|  | RMS | % of A-var | Relative to IID |
| $A$ - All | 3.437 | 100.000 | 1.000 |
| $D$ - Fast | 2.406 | 48.980 | 0.980 |
| $S$ - Slow | 2.454 | 50.948 | 1.020 |
| $E$ - Edge | 0.092 | 0.072 | 0.860 |
| $A_G$ - All Global | 0.120 | 0.122 | 274.058 |
| $D_G$ - Fast Global | 0.037 | 0.012 | 52.830 |
| $S_G$ - Slow Global | 0.114 | 0.109 | 493.227 |
| $E_G$ - Edge Global | 0.008 | <0.001 | 1,508.473 |

We also show the DSE ANOVA tables for three other subjects; HCP subject 118730 in Table S4, NYU-ABIDE subject 51050 and 51055 in Tables S5 and S6, respectively.

We observe that the cleaned data has *D*_*t*_ ≈ *S*_*t*_, which implies that the average lag-1 autocorrelation is close to zero (Sec. Appendix D.8). However, temporal autocorrelation is a ubiquitous feature of fMRI data, suggesting a contradiction. To address this, Figure 9 shows maps of the lag-1 temporal autocorrelation across the pre-processing steps. For raw data, the autocorrelation coefficient is between 0.4 and 0.6, but with successive pre-processing steps, the autocorrelation coefficient decreases until the FFP level where the median of voxel-wise autocorrelation coefficients is approximately zero. (See Fig. S18 for similar results on 20 HCP subjects).

**Figure 9:**
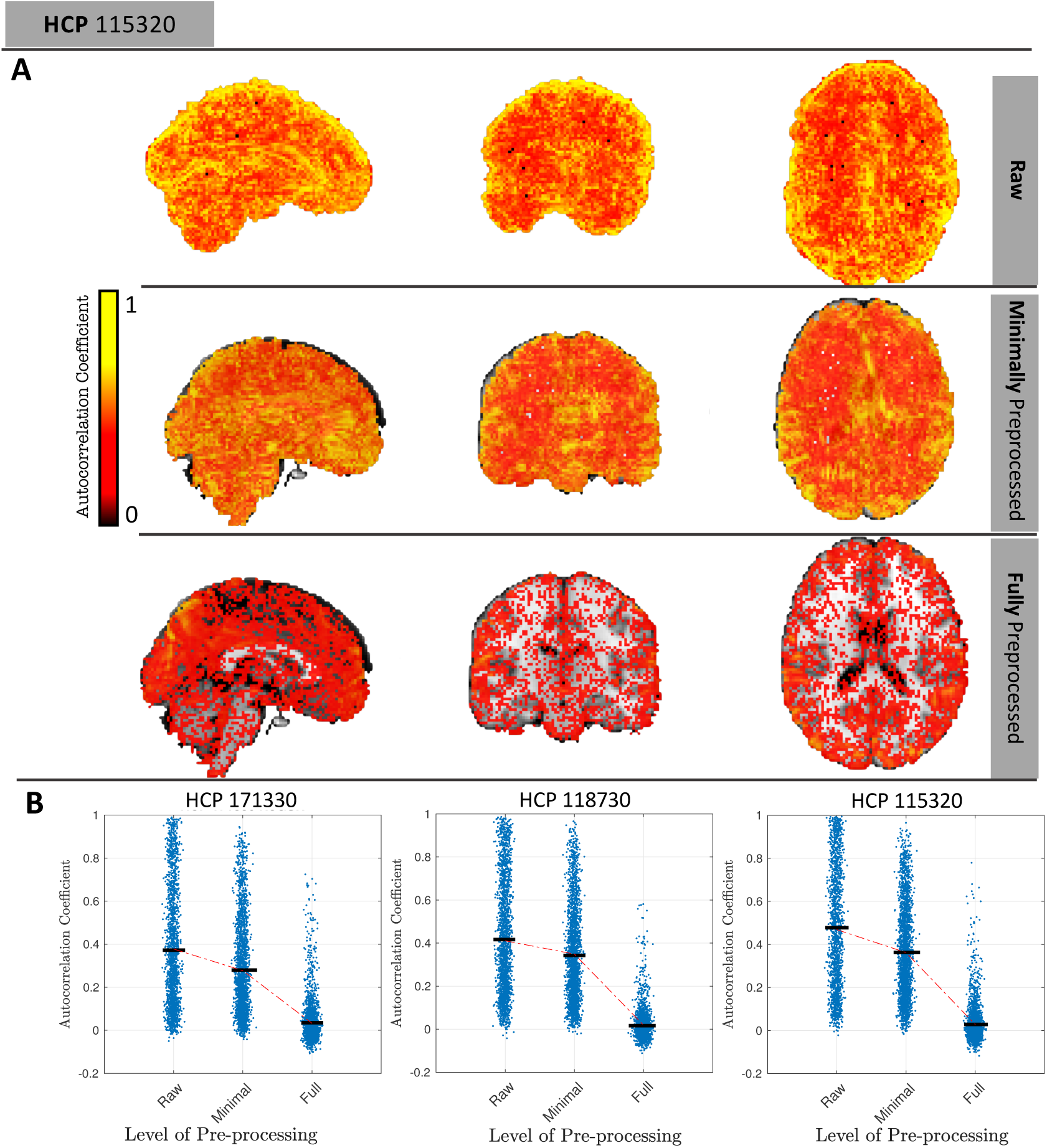
Distribution of temporal lag-1 autocorrelation across three pre-processing levels. First three rows show maps of autocorrelation for raw, minimally preprocessed and fully preprocessed, respectively, for one subject (only positive values); bottom row shows dot plots of autocorrelation for that same subject and two other subjects (random selection of 1% voxels plotted for better visualization). Fully preprocessed data has median correlation near zero, consistent with converging *S*-var and *D*-var.)

Thus, while temporal autocorrelation is present in the data, we find that the lag-1 autocorrelation coefficients do get close to zero with cleaned data, indicating that the *D*_*t*_ ≈ *S*_*t*_ heuristic is correctly indicating negligible average autocorrelation.

Figure 10 illustrates the use of the DSE decomposition to summarize the DSE components of 100 unrelated subjects in the HCP cohort, normalized as a percentage of total variance (A-var) to be maximally comparable across subjects. (See Fig. S22 for same results for ABIDE-NYU cohort). A non-normalized version of this plot (Fig. S23) is useful for viewing absolute changes, showing that S-var dramatically drops with preprocessing while *DS*-var is relatively stable.

**Figure 10:**
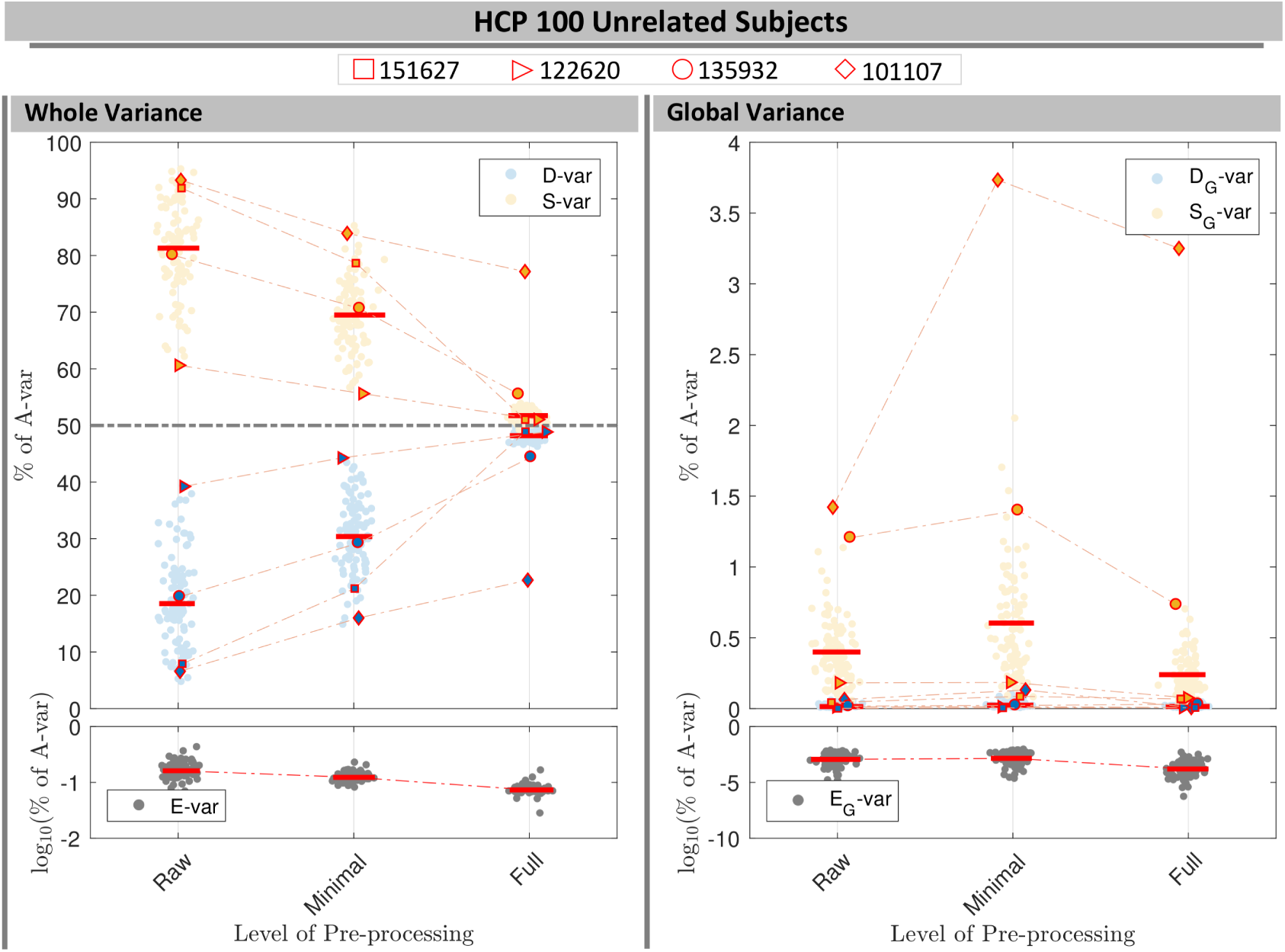
Normalized DSE decomposition for 100 HCP subjects across Raw, MPP and FPP data. The left panels show each DSE component for whole variability and the right panels illustrate the global variability of each component. Four marker types were used to follow the changes in slow and fast variability of four subjects across the pre-processing steps (see body text).

For the raw data, %*D*-var ranges from just over 5% to 40%, and *S*-var varies between 60% and 96%; the *E*-var only ever explains a negligible portion of the sum-of-squares, 0.027 to 0.50% across all three pre-processing levels. For all but two subjects the %*D*-var and %*S*-var components successively converge to 50% ±5% for FPP data.

Considering only the global variance, the slow %S_G_-var is small, usually falling well below 1%, and fast %*D*_*G*_-var is negligible, never exceeding 0.1%, reflecting the low frequency nature of the global signal.

To demonstrate the utility of the DSE decomposition in data quality control, we isolate four subjects and observe how their DSE values change with successive preprocessing.

Subject 151627, marked with a square, is one of the most extreme subjects for *S*-var and *D*-var in raw and MPP data, but has one of the smallest %*S*-var - %*D*-var differences for FPP data. This dramatic reduction in autocorrelation is confirmed in Figure 11-A, showing the cumulative distribution of lag-1 autocorrelation, and is likely linked to physiological noise around brain stem and other inferior regions (Fig. 12-A1) successfully removed by ICA-FIX (Fig. 12-A2).

**Figure 11:**
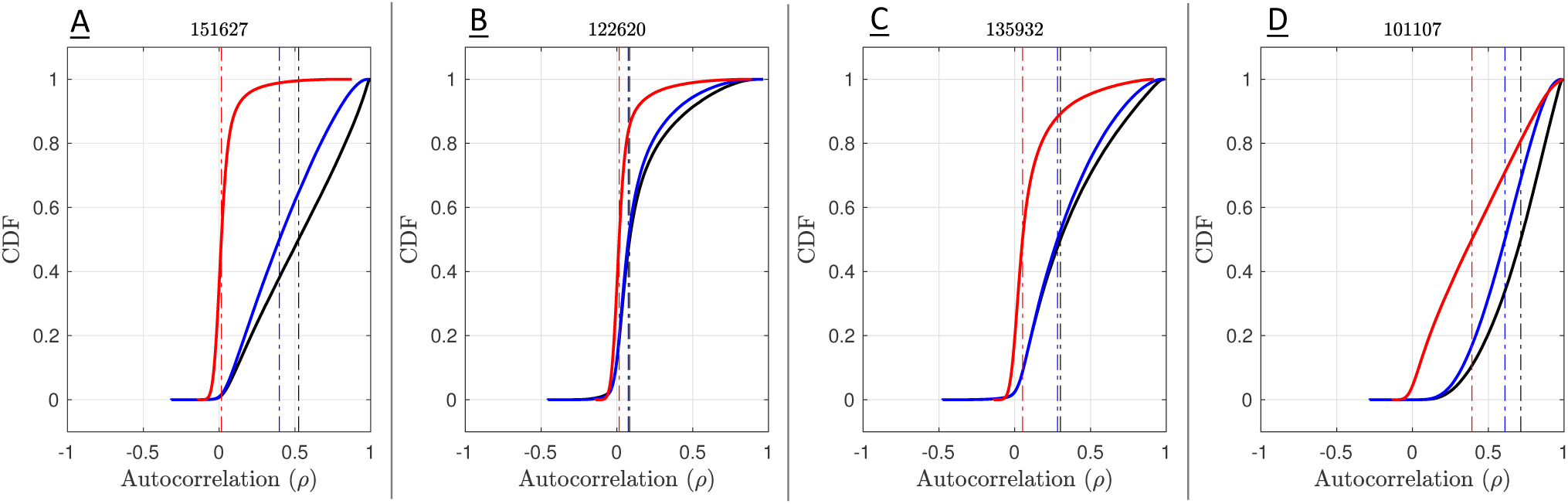
Cumulative distribution of the voxel-wise lag-1 autocorrelation coefficients for four subjects. Solid black (raw), blue (MPP) and red (FPP) lines indicates the empirical CDF and the dashed vertical lines indicate the median of autocorrelation of corresponding colors.

**Figure 12:**
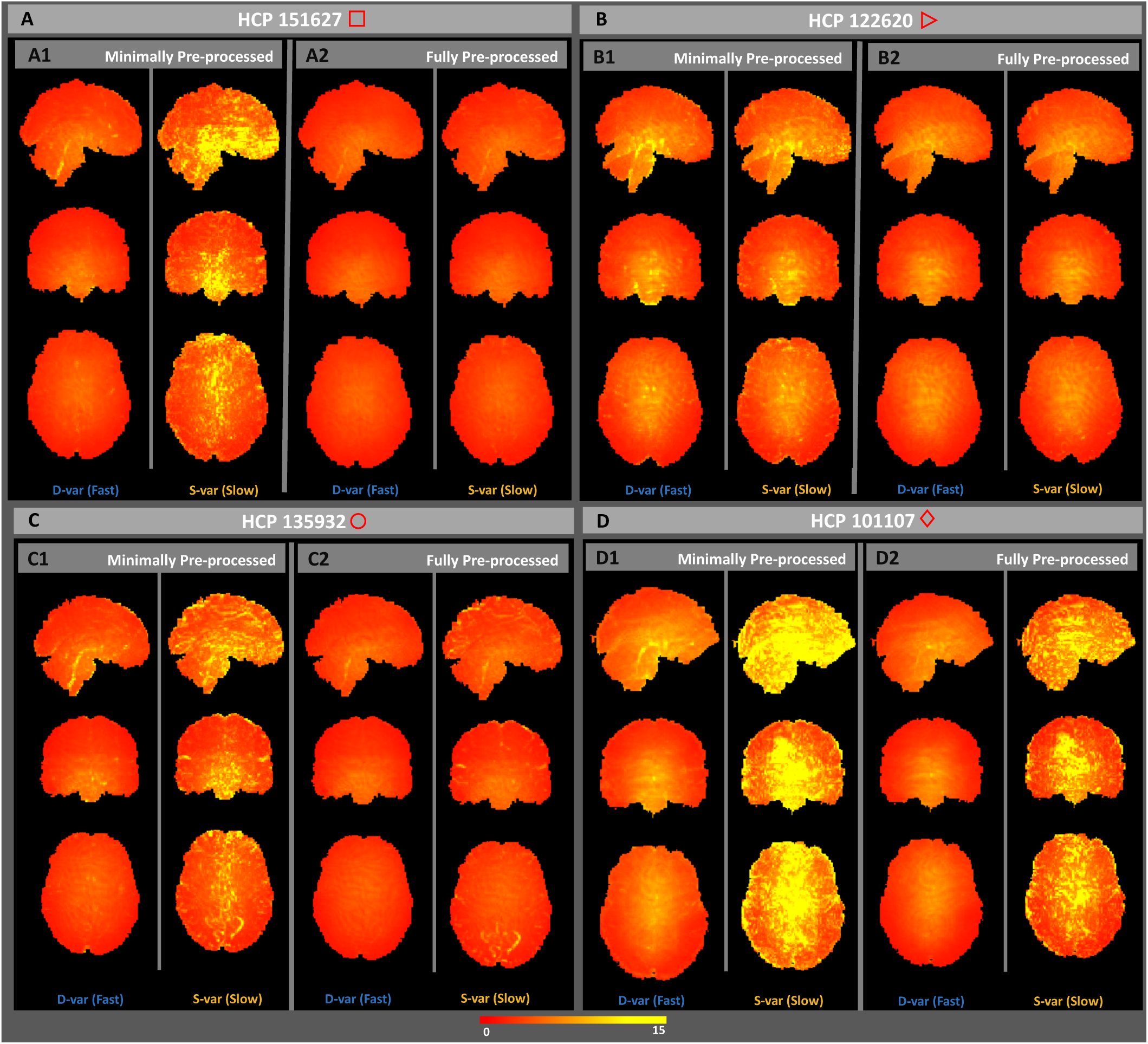
Square root *D*-var (fast) and *S*-var (slow) variability images of four subjects, for minimally (left sub-panels) and fully preprocessed data (right sub-panels). Subject 151627 appears to have been successfully cleaned, others less so; see text for detailed interpretation with respect to Figures 10 and 11

Subject 122620, marked with an triangle, has small %*S*-var - %*D*-var differences for all versions of the data, also reflected in its distribution of autocorrelation (Fig. 11-B). However, there is still some notable spatial structure in the *S*-var and *D*-var images even after clean up (Fig. 12-B2). This illustrates that if a small portion of the image possess problems, it may not be detected in any simple summary.

Subject 135932, marked with circle, has absolutely typical *S*-var and *D*-var among the 100 subjects in the raw and MPP data, but in the FPP data it has one of worst %*S*-var - %*D*-var differences. The distribution of autocorrelation coefficients reflects this (Fig. 11-C), with FPP (red line) having more large values of *ρ* than the other subjects. Inspection of the raw data *S*-var map (Fig. reffig:VarImg-C1) shows evidence of substantial structured noise that is, by in large, mostly removed by ICA-FIX correction (Fig. 12-C2). However the FPP *S*-var map shows vascular structure, likely a branch of the posterior cerebral artery near the lingual gryus; this is likely an element of physiological noise that ICA-FIX would have ideally removed but missed. Note also that this subject has low movement as measured by median FD (Fig. S21), eliminating motion as the likely source of the problem.

Finally, subject 101107, marked as a diamond, has the worst quality as measured by divergent %*S*-var and %*D*-var across preprocessing levels, with FPP level having *S*-var= 77% and *D*-var= 23%, and reflected in the largest autocorrelation values among the four subjects (Fig. 11-D). Images of *S*-var show substantial structured variability that remains even in the FPP data (Fig. 12-D), while the *D*-var image is improves notably with ICA-FIX. (This was a high-motion subject; note loss of ventromedial prefrontal cortex).

DSE time series plots of these four subjects confirm these findings, with 122620 and 151627 having flat and converged *S*-var and *D*-var time series, while 135932 and especially 101107 have structured and diverged S-var time series (Figs. S19 & S20).

To demonstrate the value of the *S*-var time series, Figure 13 explores time points where *S*_*t*_ is particularly large and small for subject 135932. Four *“S*_*t*_ images” are shown, (*Y*_*it*_ + *Y*_*i,t*+1_)^2^/4 for voxel *i,* the constituents of *S*_*t*_ (Eqn. 4). Panel A of Figure 13 shows a ‘clean’ time point, with a minimum of structured noise apparent, while panels B-D all show a similar vascular pattern. Examination of the ICA components fed into FIX finds 3 components that reflect this vascular structure that were classified as ‘good’ (Fig. S24). This demonstrates the value of the DSE decomposition to identify subtle structured noise in the data.

**Figure 13:**
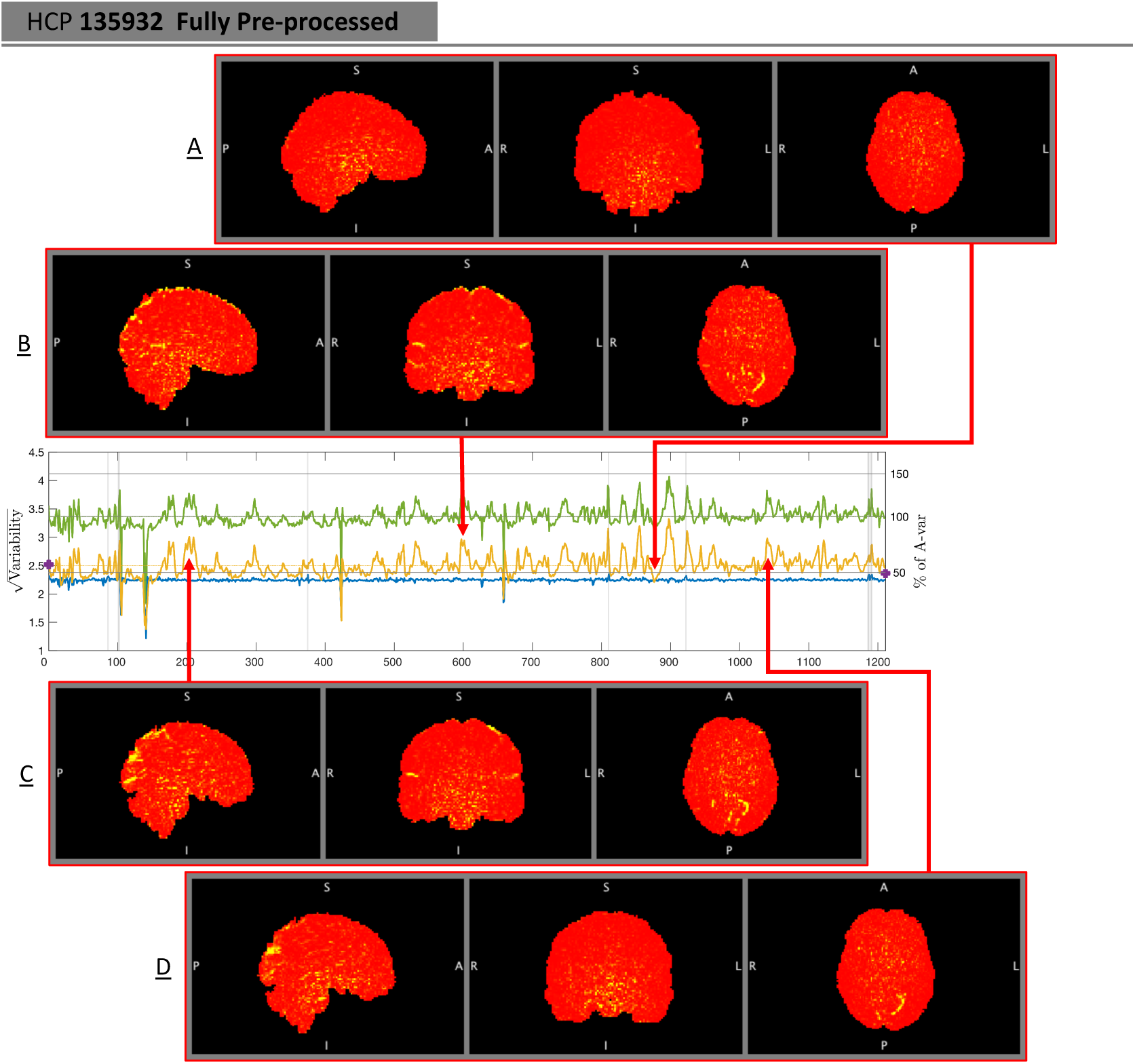
Investigation of *S-*var, slow variability artifacts. When *S*_*t*_ and *D*_*t*_ coincide, like at *t* = 871 (Panel A), the *S-*var image shows no particular structure. In contrast, we find multiple *S*-var excursions correspond to a common pattern of vascular variability across the acquisition, with time points *t* = 591, 202 and 1030 shown in panels B, C and D, respectively.

Finally, in addition to using DSE plots to investigate the quality of scans across pre-processing levels, they can also be used as a universal measure to compare the quality of scans across cohorts, data-sets and pipelines. We computed the DSE decomposition of 530 healthy subjects across 20 acquisition sites in the ABIDE dataset (Fig. S25), identifying particular sites (e.g. NYU & OHSU) and the CPAC preprocessing pipeline generally to have minimal temporal autocorrelation as reflected in *S*-var/*D*-var divergence.

## 5. Discussion

We have provided a formal context for the diagnostic measure DVARS, showing 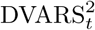 to be part of a decomposition of sum-of-squares at each successive scan pair and over the whole 4D data. We have proposed a significance test for the DVARS measure which, when detected scans are removed based on p-values, we found to address corruptions of FC while preserving temporal degrees of freedom better than other arbitrary approaches. We have also proposed the DSE decomposition which is particularly useful for summarizing data quality via DSE plots and DSE ANOVA tables. These tools concisely summarize the interplay of the fast, slow, total and global sum-of-squares, and our derived nominal expected values for each table entry facilitates the identification spatial and temporal artifacts.

Our analysis shows that D-var (and DVARS) scales with overall noise variance, and is deflated by temporal autocorrelation. We observe that as data becomes cleaner, and the background noise falls, we have greater power to identify 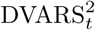 spikes. Therefore, to avoid ‘over-cleaning’ the data we complement the statistical significance of DVARS p-values with the practical significance of ∆%*D*-var, a standardized measure of the excess variance explained by a spike as a percentage of average variance. Consequently, the final candidate time-points to be scrubbed is a conjunction of statistical and practical significance; we choose a 5% familywise error rate significance level via Bonferroni and a 5% ∆%D-var cut-off; this practical significance threshold worked adequately in the HCP data we examined but may need to be recalibrated for other data sources.

Yet one more advantage of using *χ*^2^ tests, proposed in this work, is that we can estimate the effective spatial degrees of freedom which may prove to be a useful index of spatial structure in the data, but we stress this particular *χ*^2^ degrees-of-freedom *ν* is specific to this setting and is unlikely to be useful in other contexts (e.g. as a Bonferroni correction over space).

Besides ∆%*D*-var, we have introduced two standardised measures which facilitate the inter-cohort comparison of the fast (or DVARS) component regardless of intensity normalisation used in the pre-processing pipelines. For example, standardised measure %*D*-var shows the proportion of variability which can be explained via fast component while *%S*-var shows the similar proportion for the slow variability in data.

The DSE plots allow *D*-var to be judged relative to *S*-var, checking for convergence to approximately 50% of *A-*var as data approaches temporal independence, and consequently the level of autocorrelation as measure of corruption can be tightly monitored across pre-processing steps.

Using DSE plots we found two HCP subjects (101107 & 136932) where the motion-parameter regression and further ICA-FIX algorithm failed to clean the data and clearly stand out from others in the 100 unrelated subject cohort. We have used the DSE variability images to temporally and spatially locate the corruptions. It is important to note that the DSE decomposition technique should only be used before any form of resting-state bandpass filtering (such as 0.01Hz-0.1Hz) and autocorrelation modelling (such as FILM pre-whitening techniques).

Finally, we stress that we don not believe there is any one strategy can address all fMRI artifacts. Each method used in this work has it is merits and pitfalls. For example, while scrubbing was shown to be useful to remove the head motion induced spikes, it fails to remove the nuisance due to physiological signals on it is own and requires alternatives like ICA-based methods. Regardless of method, we still see value of using DSE plots and images throughout the analysis to choose a right combination of methods; see Ciric et al. (2016) for a recent comparison of various combinations of artifact methods.

### 5.1. Limitations and Future Work

Our DVARS p-values depend critically on accurate estimates of 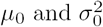. Despite finding exact expressions for the null mean and variance, we found the most practical and reliable estimates to be based on the sample 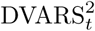 time series itself, using median for *μ*_0_ and hIQR to find *σ*_0_.Of course this indicates that our inference procedure can only infer relative to the background noise level of the data, picking out extreme values that are inconsistent with our approximating 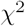 approximation.

There are two essentials avenues as continuation of this work. First is to study the effect of global signal regression via DSE decompositions. As regressing out the global mean deflates the global segment of each variability component, the DSE decomposition can be used to investigate whether global signal regression is helpful to suppress the spatial artifacts. Second, both cleaning algorithms used in this work, scrubbing and ICA-FIX, leave down-spikes (or dips) after regressing out the nuisance. These down-spikes may also affect the FC and could be detected with a two-sided variant of our hypothesis test.

## Software and Reproducibility

In this work majority of the analysis have been done on MATLAB 2015b and MATLAB 2016b, supported by FSL 5.0.9 for neuroimaging analysis.

Inference on DVARS as well as DSE decomposition techniques proposed in this paper is available via MATLAB scripts, found at http://www.github.com/asoroosh/DVARS. Also, a dedicated web page, http://sorooshafyouni.com/shiny/DSE/, present the DSE decompositions of HCP and ABIDE cohort and is regularly updated with new publicly available resting-state data sets.

Results and figure scripts presented in this work is publicly available on http://www.github.com/asoroosh/DVARS_Paper17.

## Acknowledgements

We thank Jonathan Power, Matt Glasser, Deanna Barch and Steve Peterson for feedback on the original standardized DVARS work, Javier Gonzalez-Castillo for input on interpreting our DVARS inferences, Abraham Snyder for input on history of DVARS and Michael Harms for his useful comments on the final draft of the manuscript.

Data were provided in part by the Human Connectome Project, WU-Minn Consortium (Principal Investigators: David Van Essen and Kamil Ugurbil; 1U54MH091657) funded by the 16 NIH Institutes and Centers that support the NIH Blueprint for Neuroscience Research; and by the McDonnell Center for Systems Neuroscience at Washington University.

## Appendix A. DVARS History

As far as we are aware, DVARS was first used to compute frame censoring by Smyser et al. (2011). Power et al. 2012 reported the first systematic analysis of DVARS in relation to FD in resting state fMRI. However, at least as early as 2006, a web page at the Cambridge Cognitive Brain Unit maintained by Matthew Brett’s titled “Data Diagnostics” offered tsdiffana.m, a Matlab script that produces the same measure (see http://imaging.mrc-cbu.cam.ac.uk/imaging/DataDiagnostics; when viewed on 28 October, 2012, the page listed the “last edited” data as 31 July 2006) and there are likely earlier uses in fMRI.

The idea of working with differences dates to at least 1941 in the statistics literature in work John von Neumann and colleagues (von Neumann et al., 1941). That work focused on estimation of “standard deviation from differences” when the mean slowly varied from observation to observation. They point out that the idea can traced back further, as early as 1870. In signal processing this estimator can be known as the Allan variance, developed as a robust variance estimator in the presence of 1/f noise (Allan, 1966). In cardiology the “root mean square successive difference” is a standard measure of heart period variability (Berntson et al., 2005), and as “mean successive squared difference” (MSSD) it has recently been used in neuroimaging as an index neuronal variability (Samanez-Larkin et al., 2010; Garrett et al., 2013). For yet more background see Kotz et al. (1988).

Despite successive work on finding the exact distribution of this variance estimate (Harper, 1967), or using it in a test for the presence of autocorrelation (Cochrane and Orcutt, 1949), we are unaware of any study of the distribution of the individual differences averaged over a multivariate observation, as is the case in this fMRI application.

## Appendix B. Plotting the global variance decomposition

The global variance components, at each time point, are just a single scalar value squared. Thus they may be more intuitively plotted in a signed RMS form. For example, instead of plotting variance *A*_*Gt*_, *D*_*Gt*_ and *S*_*Gt*_, the signed quantities

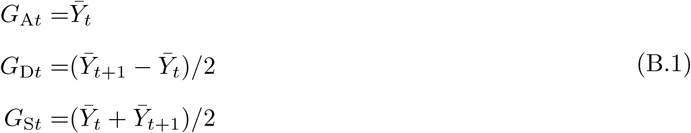

can be plotted. These set of three time series may seem arbitrary, but have the feature of the sum of squares of *G*_*Dt*_ and *G*_*St*_ sum to the mean-square *G*_*At*_ and *G*_A*t,t*+1_.

## Appendix C. Derivation of DSE variance decomposition

The decomposition of the average variance at time *t* and *t* + 1, Eqn. (5), is based on a simple algebraic identity; for variables *a* and *b*,

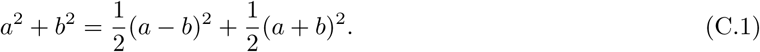

This justifies a decomposition of the average variance at each voxel i, for each time *t* = 1,…, *T* - 1,

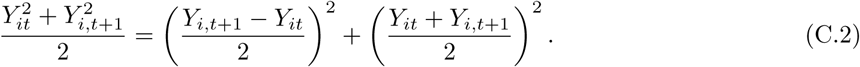

Averaging this expression over voxels *i* = 1,…, *I* gives the decomposition for scan pair variance A_*t,t*+1_ in Eqn. (5). Summing image variance A_*t,t*+1_ over *t*, however,

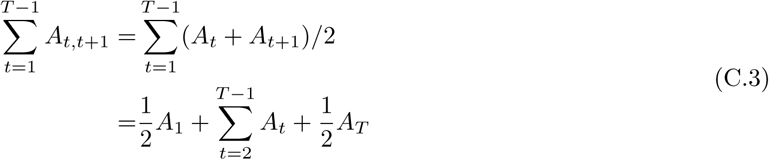

misses 1/2 of edge terms, which are added to produce the fundamental DSE decomposition in Eqn. (7).

## Appendix D. Derivation of DSE ANOVA Mean Squares

Here we set out the least restrictive model possible to justify our expected values for the DSE ANOVA table (Table 1). While the DSE ANOVA table and decompositions *A = D + S + E* and *A*_*G*_ = *D*_*G*_ + *S*_*G*_ + *E*_*G*_ are in mean-square (MS) units, below we develop the results in terms of sum-of-squares (SS) that, in each case, can be divided by *I* × *T* to obtain the MS.

All of the results follow from application of rules for expectations and variances of quadratic forms of mean zero vectors. For reference, if *w* is a mean zero random vector with covariance Σ, and *B* is a square matrix, then 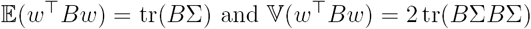.

### Appendix D.1. Model

In defining the the joint distribution of all *I* × *T* elements of the 4D data {*Y*_*it*_}, we will always assume is that *Y*_*it*_ is mean zero and has constant variance over time, 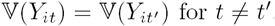, but allow variance to vary over space. For data organized as time series, length-*T* vectors *Y*_*i*_, let

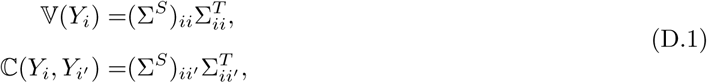

where Σ^*S*^ is the *I × I* spatial covariance matrix, common to all time points, and (Σ^*S*^)_*ii*_ is the variance at the ith voxel, 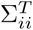 is the *T × T* temporal autocorrelation matrix for voxel *i,* 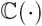 denotes covariance, and 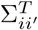, is the *T × T* temporal cross correlation matrix for voxels *i* and *i*′. This implies that, for data organized as images, length-*I* vectors *Y_t_,*

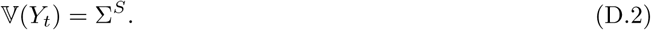

When a time-space separable covariance structure is assumed then 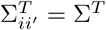 for all *i*, *i*'.

### Appendix D.2. A-var Expected SS

Total SS 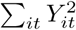 has expected value

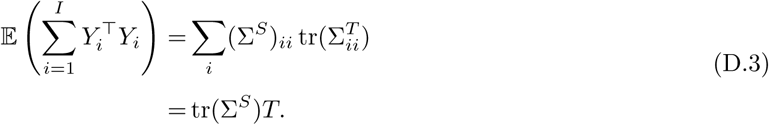

### Appendix D.3. D-var and S-var Expected SS

The total *D*-var SS is 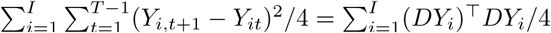 where

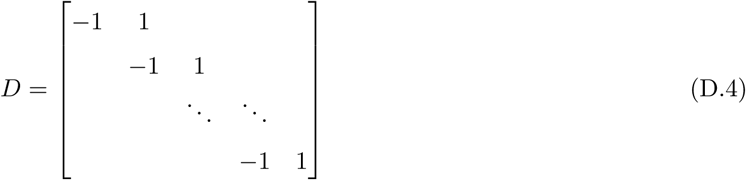

is the (*T -* 1) *× T* finite difference matrix. We have

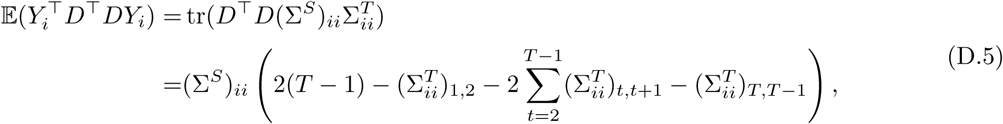

where notably the last expression only depends on the lag-1 temporal autocorrelations. To obtain more interpretable results we further assume that there is a constant lag-1 autocorrelation at each voxel, 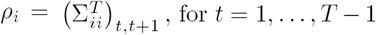, which reduces (D.5) to 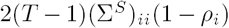. This gives the expected total *D*-var SS as

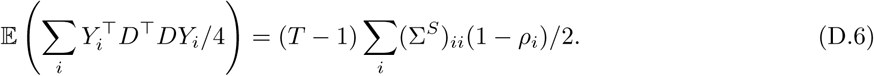

If we yet further assume constant temporal autocorrelation *ρ,* corresponding to our separable model, this SS simplifies to 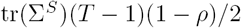.

The expected SS for *S*-var is follows the same arguments with differencing matrix replaced with a running sum matrix abs(*D*), negating the three negative terms in Eqn. D.5, and reducing to 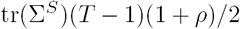 under spatially and temporally homogeneous lag-1 temporal autocorrelation.

### Appendix D.4. E-var Expected SS

The total SS E-var is 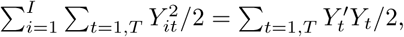, with expected value

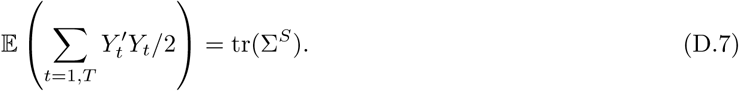

### Appendix D.5. A_G_-var Expected SS

The global time series is 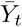 and total SS due to global is

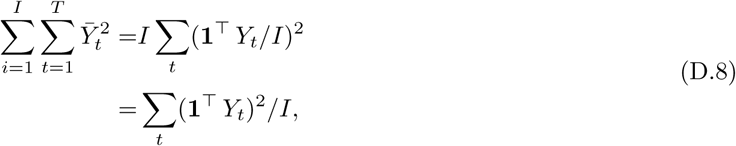

where **1** is a vector of ones. The expectation of the squared term is 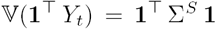, and thus the expected SS is

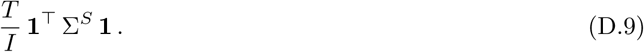

### Appendix D.6. D_G_-var and S_G_-var Expected SS

Write the global differenced time series as 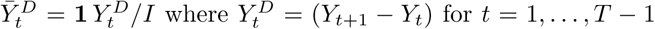.

The total SS due to half differenced global *D*_*Gt*_ is then

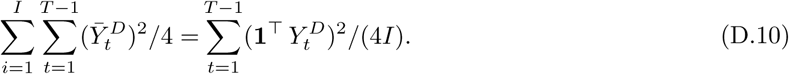

To find the expectation of the squared term, note that

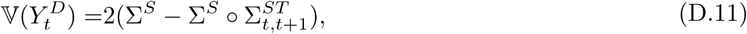

where ° is the Hadamard product and 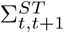 is the spatiotemporal covariance matrix, elements extracted from the temporal cross correlation matrix as per 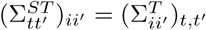, and that

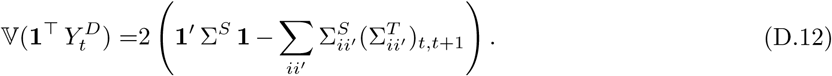

The final expression for the expected SS is then, with successive assumptions

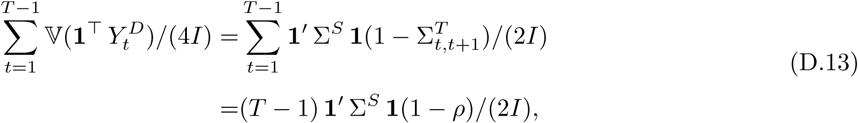

where first equality comes from assuming a separable covariance structure and the second from a common lag-1 autocorrelation.

The result for *S*_*G*_-var follows similarly.

### Appendix D.7. EG-var Expected SS

The total SS *E*_*G*_-var is 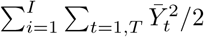, and following same arguments as for *A*_*G*_-var has expected value

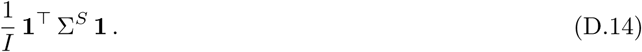

Results for the non-global terms in the decomposition *A*_*N*_ = *D*_*N*_ + *S*_*N*_ + *E*_*N*_ follow as difference of respective total and global terms.

### Appendix D.8. Expected value of the difference of percent S-var & D-var

The convergence of *S*-var and *D*-var is a visual diagnostic indicating cleaned data. Here we find the expression for the difference of the average normalized *S*-var and *D*-var measures. The most general case is found using Equation D.5:

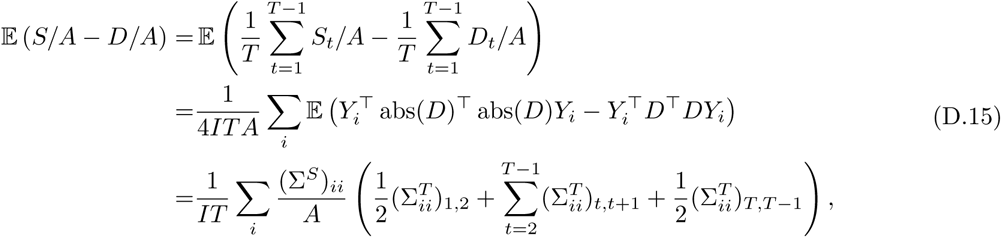

where we’ve assumed *A* has negligible variability. This result can be seen to be a variance-weighted average of lag-1 temporal autocorrelations over time and space. It can also be shown that a similar result holds for each time *t* = 1,…, *T* - 1,

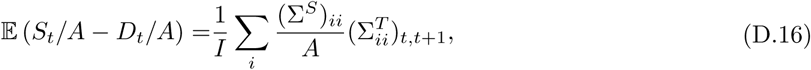

If we assume 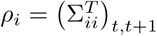, i.e. time-constant lag-1 autocorrelations at each voxel, D.15 reduces to

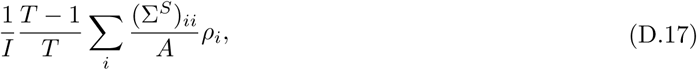

as does D.16 but without the (*T* - 1)/*T* term.

These results show that the difference between normalized *S*-var and *D*-var is a weighted average of lag-1 autocorrelations.

## Appendix E. Derivation of DVARS Null Distribution

As results are more naturally defined for squared quantities, we seek a null distribution for

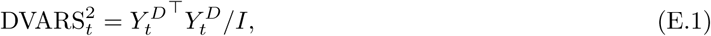

where 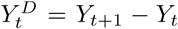 as above. While an expression of the mean of DVARS can be obtained from Eqn. (D.11), note also

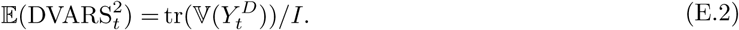

That is, the expected value of 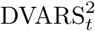 is simply the variance of each voxel in the differenced data, averaged over voxels. The natural estimator of this is the sample mean (or robust equivalent) of the sample variance image (or robust equivalent) of the differenced 4D data.

The variance is more involved

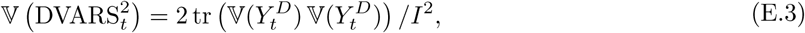

in particular depending on the entirety of the *I* × *I* difference image variance matrix. For the most restrictive assumptions considered above 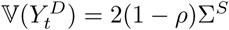 and thus

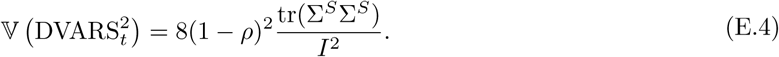

This dependence on the full spatial covariance demands the empirical approaches to variance estimations taken in the body of the paper.

Only at this point do we invoke a normality assumption, and make use of the classic chi-square approximation for sums-of-squared normal variates (Satterthwaite, 1946). In this approach we equate the mean and variance of 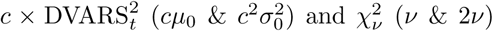 and solve for *c* and *ν,* giving the multiplier 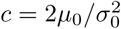 and degrees-of-freedom 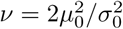 as found in Section 2.4.

## Appendix F. Power Transformations to Improve DVARS Variance Estimation

The robust IQR-based variance estimate reflects a normality assumption, equating the sample IQR with that of a standard normal. 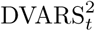, as a sum-of-squares and as reflected by its *χ*^2^ approximation, may exhibit positive skew. Hence we consider power transformations of 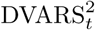 that may improve symmetry and the accuracy of the IQR variance estimate. While the asymptotically optimal power transformation to normality for *χ*^2^ is known to be the *d =* 3 cube-root transformation (Hernandez and Johnson, 1980), our test statistic is only approximately *χ*^2^ and, in particular, variance heterogeneity can worsen the approximation.

To obtain a quantity that should be more symmetric consider the power transformation

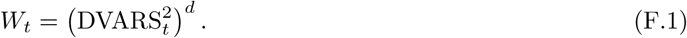

IQR-based estimates of the variance of *W,* 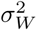, will hopefully be more accurate than such estimates on DVARS^2^. However, ultimately we seek estimates of the variance of DVARS^2^, and so for a given *d* we compute

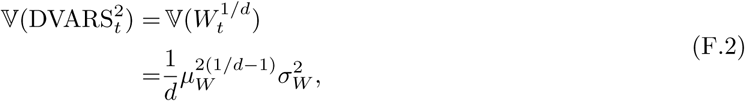

where the last expression is the delta method variance of 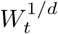, and *µ*_*W*_ is the mean of *W*_*t*_ (which we robustly estimate with the median of *W*_*t*_).

## Footnotes

1 Going forward we drop the third row of the DSE ANOVA table showing non-global variance, since in practice the global explains so little variance that the first and third rows are essentially the same; see e.g. Table 7 entries’ for *A_G_,* and Fig. 10 right.

